# Intercellular Transfer of PTBP1 Drives Human Neural Stem Cell Fate

**DOI:** 10.64898/2026.02.19.706765

**Authors:** D.L. Capobianco, F. Di Palma, E. Filomena, C. Lasconi, C. Pousis, L. Simeone, F. Proto, D.C. Profico, M. Gelati, E. Picardi, G. Pesole, A.L. Vescovi, M. Svelto, L. Simone, A.M. D’Erchia, F. Pisani

## Abstract

During fetal brain development, temporally defined alternative splicing (AS) programs control human neural stem cell (hNSC) self-renewal and differentiation, thereby regulating neurogenesis and gliogenesis. Polypyrimidine tract-binding protein 1 (PTBP1) is a master regulator of AS during neurogenesis; however, its functional role and dynamics in hNSCs remain largely unexplored.

Here, we investigate the cellular and molecular functions, nucleocytoplasmic distribution, and intercellular trafficking of PTBP1 in primary hNSCs.

We found that PTBP1 knockdown (KD) alters self-renewal capacity, mitochondrial dynamics and membrane potential, lipid droplet abundance, and PTBP2 expression.

RNA sequencing analysis revealed that PTBP1 depletion affects the expression profiling of hundreds of coding and non-coding genes, collectively orchestrating a neuronal differentiation program.

Super-resolution τ-STED microscopy and live-cell imaging demonstrated that PTBP1 localizes not only to the nucleus but also to the cytoplasm, tunneling nanotubes (TNTs), migrasomes, and extracellular vesicles (EVs). Co-culture experiments and EV uptake assays showed that cytosolic PTBP1 can be transferred between hNSCs and delivered to the nuclei of recipient cells via TNTs and EVs. Moreover, EVs were found to contain specific and previously uncharacterized PTBP1 isoforms and were efficiently transferred to PTBP1-KD cells, rescuing their proliferative capacity. Analysis of the mouse brain reveals the presence of PTBP1 in the V-SVZ and within TNT-like structures connecting NSCs, suggesting a role for TNT-mediated PTBP1 trafficking in vivo.

Together, these findings uncover previously unrecognized roles for PTBP1 in hNSCs and provide the first evidence that PTBP1 can be transferred between hNSCs via TNTs and EVs, revealing a novel mechanism by which hNSCs may regulate fetal neurogenesis.

**Graphical Abstract:** **A:** PTBP1 regulates hNSC fate by controlling cell proliferation, lipid droplet dynamics, mitochondrial function, and post-transcriptional programs involved in neuronal commitment.

**B:** Cytosolic PTBP1 is transferred between hNSCs via tunneling nanotubes (TNTs) and extracellular vesicles (EVs).

Abbreviations: LV, lateral ventricle; aRG, apical radial glial cells; bRG, basal radial glial cells; V-SVZ, ventricular–subventricular zone; EVs, extracellular vesicles; TNTs, tunneling nanotubes.

## Introduction

During fetal brain development, apical and basal radial glial cells (aRGs and bRGs)—collectively referred to as neural stem cells (NSCs)—sequentially differentiate into neurons, astrocytes, and oligodendrocytes through precise spatiotemporal coordination of NSC self-renewal and lineage specification ^1–4^

Most known mechanisms regulating NSCs rely on paracrine signaling or cerebrospinal fluid–derived cues, including fibroblast growth factors (FGFs), insulin-like growth factors (IGFs), Sonic hedgehog (Shh), retinoic acid, bone morphogenetic proteins (BMPs), and Wnts—molecules essential for aRG survival, proliferation, and neurogenic potential ^5–7^. In this context, the spatiotemporal control of alternative splicing (AS) has emerged as a key regulatory mechanism. AS modulates NSC fate at multiple levels by controlling enzymes and signaling pathways critical for self-renewal and differentiation, including the Wnt, FGF/ERK, and NOTCH pathways, among others ^8^.

Polypyrimidine tract-binding protein 1 (PTBP1) is a well-characterized AS-regulator that has been largely investigated for its role in controlling the fate of NSCs and embryonic stem cells in mouse through multiple NSC-related pathways ^9–15^. PTBP1 is primarily known as a nuclear protein acting as splicing regulator but, interestingly it can shuttle between the nucleus and cytoplasm ^16^, where it regulates mRNA stability and translation ^17,18^. PTBP1 has also been also extensively investigated as a target for astrocyte-to-neuron conversion strategies in the treatment of neurodegenerative diseases ^19,20^. However, its subcellular localization, dynamic and functional role in primary non-immortalized human NSCs (hNSCs) from the sub-ventricular zone remain unknown.

Tunneling nanotubes (TNTs) are actin-based structures that mediate the intercellular transfer of organelles, ions, proteins and other cytoplasmic components ^21,22^ and are increasingly recognized as important mediators of neural communication ^23–33^, including in vivo, as recently demonstrated for inter-dendritic TNTs in the mouse brain ^34^. We have previously identified TNTs hNSCs and demonstrated active organelle exchange through these structures ^32^. However, whether cytosolic form of PTBP1 exist in hNSCs and can be trafficked through TNTs remains unknown.

TNTs are not the only means by which cells can exchange cytosolic proteins; extracellular vesicles (EVs) can also mediate this type of trafficking. In particular, among the different EV subtypes, microvesicles—typically ranging from 50 nm to 5 µm and formed by outward budding and fission of the plasma membrane ^35^—can facilitate the intercellular transfer of cytosolic components. Other EV subtypes, such as migrasomes, which are derived from the retraction fibers of migrating cells, also contribute to intercellular communication by exchanging cytosolic components ^36,37^

Numerous studies have examined the content and functions of EVs derived from NSCs, particularly focusing on the therapeutic applications of exosomes and their capacity to influence differentiation in recipient cells ^38–40^. For example, exosomes derived from rat SVZ NSCs can direct neural lineage specification in human mesenchymal stem cells ^41^, while exosomes secreted by immature neurons promote neuronal differentiation of mouse NSCs in vitro ^42^. However, the role of hNSC-derived EVs in NSC-to-NSC communication and in the regulation of self-renewal remains largely unexplored, and whether PTBP1 can be trafficked between hNSCs via EVs is still unknown.

Here, we investigated the cellular and molecular roles, subcellular localization, and dynamics of PTBP1 in hNSC fate and tested the hypothesis that TNTs and EVs mediate the transfer of cytosolic PTBP1 between hNSCs, thereby contributing to the spatial regulation of NSC fate.

We found that PTBP1 regulates hNSC fate at multiple levels, including cell proliferation mitochondria, lipid droplets and differentiation, controlling the transcriptional profile of hundreds of coding and non-coding RNA. Notably, we demonstrated that PTBP1 localizes to the nucleus, cytoplasm, TNTs, migrasomes, and EVs in an isoform-dependent manner.

Moreover, we demonstrated that PTBP1 is transferred between hNSCs via TNTs and extracellular vesicles (EVs), enters the nuclei of PTBP1-depleted recipient cells, and contributes to the restoration of cell proliferation. We also provide the first evidence of PTBP1-containing TNT-like structures between NSCs in the mouse SVZ.

Together, our findings provide the first evidence that TNTs and EVs mediate the intercellular transfer of PTBP1 between NSCs, suggesting that this mechanism may play a role in the spatiotemporal regulation of brain development.

## Methods

### Human neural stem cell isolation and culture

Human brain tissue (GW16) was immediately transferred under strict sterile conditions to the GMP facility in a controlled environment. Then brain specimen was washed in a PBS solution (Dulbecco’s PBS 1X, Carlo Erba Reagent) supplemented with 50 μg/ml of gentamicin and dissociated to reach a monocellular suspension mechanically. Cells were seeded at a density of 10^4^ cells/cm^2^ in a chemically defined culture medium^43^ with EGF 20 ng/ml and bFGF 10 ng/ml. Cultures were maintained in a humidified hypoxic incubator at 37°C, 5% O_2_, and 5% CO_2_ and cells were allowed to proliferate as free-floating neurospheres. Approximately 7–10 days after the primary cell seeding, neurospheres were collected in 15 mL tube, centrifuged, the supernatant discharged, and the cell pellet mechanically dissociated using p200 micropipette. The obtained single-cell suspension was stained with 0.4% Trypan Blue stain solution (Invitrogen, cat #T10282) and counted using a Burker chamber. A cell viability near 95–98% was measured in each neurospheres passage. Throughout these passages, aliquots of cells were frozen as neurospheres and cryopreserved in a culture medium with 10% dimethyl sulfoxide. When adhesion culture was necessary glass supports were coated with Cultrex (BME, Pathclear cat #3432-005-01) 1:50 in hNSC-medium ^44^ for 1 h at 37 °C in according to manual instruction.

### PTBP1 knockdown

hNSCs were either plated on Cultrex™-coated cover glasses at a density of 1.6 × 10^4^ cells/cm² or treated in suspension at the same seeding density. Cells were exposed to Accell Human PTBP1 (5725) siRNA - SMARTpool (Dharmacon™, #E-003528-00-0020) and Accell Non-targeting Control Pool (Dharmacon™, #D-001910-10-20) at a final concentration of 1 µM directly in the culture medium. Treatments were maintained for 7–10 days.

### Lentiviral constructs

The coding sequence of PTBP1 isoform a (NM_002819.5), fused at the C-terminus to the mCherry coding sequence, was cloned into the lentiviral expression vector pLV[Exp]-Puro-CMV. This construct was transfected into packaging cells, and the resulting RNA-based lentiviral particles were subsequently produced, isolated, and titrated. A lentivirus encoding mCherry alone was used as a negative control. All procedures were performed as a service by VectorBuilder.

### Lentiviral infection

For overexpression experiments, hNSCs in suspension were infected with lentiviral (LV) particles expressing PTBP1 isoform-a (NM_002819.5) (PTBP1-mCherry-LV) or the corresponding control virus expressing mCherry alone (mCherry-LV) using Multiplicity of infection (MOI) = 4.

Twenty-four hours post-infection, the culture medium was replaced, and cells were washed three times to remove residual virus. This was achieved by centrifuging the cells at 200 × g for 10 minutes, carefully aspirating the supernatant, and resuspending the cell pellet in fresh culture medium. Cells were then returned to standard culture conditions for continued growth and expression analysis.

### Immunofluorescence

To minimize breakage of TNTs, hNSCs fixation with 4% paraformaldehyde was performed for 10 min at room temperature by gently replacing the culture medium. Samples were then carefully washed with PBS, permeabilized with 0.3% Triton X-100 in PBS for 15 min, and blocked with 3% BSA in PBS for 30 min. Primary antibodies were incubated in 3% BSA in PBS for 24 h at 4 °C. In some experiments, to maximize primary antibody yield, incubation was performed overnight in 0.3% Triton X-100/3% BSA in PBS. After primary antibody incubation, samples were gently washed multiple times with 3% BSA in PBS and then incubated with secondary antibodies for 1 h at room temperature in 3% BSA in PBS. Finally, samples were extensively washed and mounted in Vectashield antifade mounting medium (Vector Laboratories, cat. #H-1000-10).

### Staining of mitochondria for fixed-cell-based microscopy analysis

To distinguish recipient cells from PTBP1-mCherry donor cells, in 24h co-culture experiments the mitochondria of recipient hNSCs were labelled prior to coculture. Briefly, a single-cell suspension of recipient hNSCs was incubated with 100 nM MitoTracker Deep Red FM (Invitrogen, cat. #M22426) diluted in hNSC medium for 30 min. After incubation, cells were centrifuged and resuspended in fresh prewarmed hNSC medium at least twice, then left overnight in hNSC medium and centrifuged again. The cell pellet was washed once to completely remove excess dye, and cells were seeded onto Cultrex-coated glass coverslips.

Subsequently, PTBP1-mCherry–expressing donor hNSCs were added. After 24 h of coculture, cells were fixed with 4% paraformaldehyde, as previously described, and processed for immunofluorescence according to the protocols described above. In coculture experiments, PTBP1-mCherry protein was identified using an anti-mCherry antibody [EPR20579] Rabbit mAb (Abcam, cat. #ab213511) at a 1:500 dilution.

### Confocal and Tau-STED microscopy

All image acquisitions were performed in serial acquisition mode between frames. XYZ image stacks were acquired with a raster size of 1024 × 1024 pixels in the X–Y plane and a z-step of 0.2 μm between consecutive optical sections. Imaging was performed using a Leica Stellaris 8 microscope operated in confocal mode or, when indicated, in Lightning or Tau-STED super-resolution mode.

Fluorescence excitation was provided by a white light laser, and emission signals were detected using hybrid detectors. Tau-STED acquisition were applied only when specified. Three-dimensional (3D) reconstructions and projections from z-stacks were generated using Leica Application Suite X (LAS X). Image visualization and quantitative analyses were performed using LAS X and FIJI software using 3D-objects counter plugin.

### Time-lapse live-cell holotomography/fluorescence imaging and holotomographic analysis of lipid droplets and mitochondria under physiological hypoxia

Live-cell 3D imaging was performed using a Nanolive 3D Cell Explorer Fluo (Holotomographic Microscope CX-F) combined with a CX-F Incubator Hypoxia Module (Okolab), which maintained controlled physiological hypoxic conditions for hNSCs (5% O₂, 5% CO₂). Single cells or neurospheres were seeded on Cultrex coated-µ-Dish 35 mm low glass bottom dishes (Ibidi, cat. #80137). Images were computationally reconstructed into 3D time-lapse maps, enabling visualization and quantification of organelles, cellular dry mass, and dynamic processes such as membrane remodeling, TNTs, migrasomes, organelle organization, and lipid droplet dynamics without phototoxicity or chemical labeling, thereby preserving physiological hypoxia. Quantitative analyses were performed using Nanolive’s AI-based digital analysis suite (CX-F Analytics Module – EVE Explorer) and Fiji software. To visualize the dynamics of PTBP1-mCherry in live-cell time-lapse experiments, cells were imaged under the hypoxic conditions described above, tracking mCherry fluorescence over time using the LED-based excitation module of the CX-F system. Excitation power was kept at a minimum to reduce photodose and preserve cell viability.

### Extracellular vesicles isolation

Extracellular vesicles were isolated as previously described ^45^. After 7 days in culture, neurospheres were collected and pelleted by centrifugation at 200 × g for 10 min. The supernatant was carefully recovered and kept on ice, while neurospheres were mechanically dissociated by gentle pipetting using a P200 pipette until a single-cell suspension was obtained. The previously collected supernatant was then added back to the cell suspension.

The resulting suspension was centrifuged at 200 × g for 10 min at 4 °C to remove intact cells. The supernatant was carefully transferred to a new 50-ml Falcon tube and subjected to an additional centrifugation step at 300 × g for 10 min at 4 °C to eliminate residual cellular debris. The clarified supernatant was again recovered and transferred to a clean tube.

To isolate extracellular vesicle fractions, the supernatant was further centrifuged using a swinging-bucket rotor at 2,000 × g for 20 min at 4 °C. The resulting pellet corresponded to the 2,000 × g extracellular vesicle fraction (EV2,000g). The supernatant depleted of EV2,000g was subsequently transferred to a new tube and centrifuged using a fixed-angle rotor at 10,000 × g for 40 min at 4 °C, yielding the 10,000 × g extracellular vesicle fraction (EV10,000g).

For vesicle-mediated transfer experiments, EV2,000g and EV10,000g pellets were directly resuspended in culture medium and seeded into the appropriate wells.

For Western blot analysis, each EVs pellet was subjected to two washing steps in ice-cold PBS to remove residual culture medium contaminants. Specifically, the EV2000g pellet was washed twice by centrifugation at 2,000 × g for 20 min at 4 °C, replacing the PBS at each step, while the EV10000g pellet was washed twice at 10,000 × g for 40 min at 4 °C. Cell pellet was washed twice with ice-cold PBS and subsequently lysed directly in 4× Laemmli Sample Buffer (Bio-Rad, cat. #1610747) supplemented with 50 mM dithiothreitol (Bio-Rad, cat. #1610611) and 20 mM protease inhibitor cocktail (Roche cOmplete™, cat. #04693124001). Lysis was performed adding 4x Laemmli buffer 1:1 ratio relative to the residual PBS volume, obtaining 2x Laemmli buffer as final concentration. Similarly, EV2000g and EV10000g pellets were lysed in the same supplemented 4× Laemmli buffer using an identical 1:1 ratio.

All centrifugation and washing procedures were performed with samples kept on ice to prevent extracellular vesicle aggregation. The washing steps were essential to remove the high concentration of BSA present in the culture medium, which could otherwise interfere with Western blot analyses.

### Wide-field fluorescence imaging

Wide-field fluorescence imaging was performed using a Nikon ECLIPSE Ti-S inverted microscope equipped with 10×, 20×, and 40× dry objectives. Images were acquired and analyzed using NIS-Elements and FIJI/ImageJ software.

### Analysis of mitochondria membrane potential

Mitochondrial membrane potential was assessed using the potentiometric dye JC-1 (Invitrogen, cat. #T3168). Prior to staining, JC-1 was diluted 1:500 in prewarmed culture medium and thoroughly vortexed, as the dye is poorly soluble. The staining solution was then used to replace the existing culture medium in wells containing the cells, followed by incubation for 20 min in a humidified cell culture incubator under the conditions described above.

JC-1 accumulates in mitochondria in a membrane potential–dependent manner. At low mitochondrial membrane potential, the dye remains in its monomeric form and emits green fluorescence, whereas at high membrane potential it forms J-aggregates that emit red fluorescence. After incubation, cells were washed three times with fresh culture medium and immediately imaged to preserve membrane potential–dependent fluorescence signals.

Wide-field fluorescence imaging was performed using a Nikon ECLIPSE Ti-S inverted research microscope equipped with 10×, 20×, and 40× dry objectives. Images were acquired using NIS-Elements software and subsequently analyzed in FIJI/ImageJ.

Quantitative analysis was carried out in FIJI/ImageJ by measuring the mean fluorescence intensity in both the green and red channels. Mitochondrial membrane potential was expressed as the ratio of red to green fluorescence intensity, providing a ratiometric and concentration-independent measure of mitochondrial polarization based on the relative abundance of JC-1 aggregates versus monomers.

### Western Blotting

Total proteins dissolved in 2x Laemmli Sample Buffer (Bio-Rad, cat. #1610747), 50 mM dithiothreitol (Bio-Rad, cat. #1610611) and 20mM protease inhibitor (Roche cOmplete™, cat. # 04693124001) heated to 95°C for 10 min, resolved on a 10 or 12% Mini-PROTEAN® TGX Stain-Free™ Protein Gels (Bio-Rad, cat. #4568036), and transferred onto 0.2 µm PVDF membranes (Bio-Rad, cat. #1704156). After transfer, the membranes were blocked for 5 min with Every Blot Blocking Buffer (Bio-Rad, cat. # #12010020) and incubated with antibodies. Reactive proteins were revealed with Clarity and Clarity Max ECL Western Blotting Substrates (Bio-Rad, cat. # 1705061, cat. # 1705062) and visualized on a VersaDoc imaging system (Bio-Rad). Images were analyzed using Image Lab Software (Biorad).

### RNAseq analysis

Total RNA was extracted from PTBP-siRNA and scramble-siRNA treated hNSCs using the RNeasy micro kit (Qiagen, cat. # 74004), according to the manufacturer’s instructions. RNA was quantified on NanoDrop 2000c (ThermoFisher Scientific) and qualitatively evaluated on Bioanalyzer 2100, using the RNA 6000 Pico Kit (Agilent, cat. # 5067-1513). Three replicates for each condition were processed. RNA-seq libraries were prepared from 200 ng of total RNA, using the Illumina Stranded Total RNA Prep with Ribo-Zero Plus kit (Illumina, cat. # 20040529), according to the manufacturer’s protocol. Final libraries were checked on the Bioanalyzer 2100, using the DNA High Sensitivity Kit (Agilent, cat. # 5067-1513) and quantified by fluorimetry using the Qubit dsDNA Assay Kit (Thermo Fisher Scientific, cat. # Q32854). Sequencing was performed on Illumina Novaseq 6000 platform, generating an average of 100 millions of 100pb paired-end reads for sample. All computations were performed on machines running GNU+Linux (3.10.0–862.14.4.el7.x86_64), by using R (version 3.6.1) and Bash (4.2.46(2)-release x86_64-redhat-linux-gnu). The quality of the RNA-seq reads was preliminarily inspected with fastQC (https://www.bioinformatics.babraham.ac.uk/projects/fastqc/) and MultiQC ^46^ and trimming was performed with fastp ^47^ to remove Illumina sequencing adapters, sequences with a phred33 quality score inferior to 25, low complexity sequences, polyX sequences at 3’ ends and eventually discard reads less than 50bp long. All steps of the analysis dependent on genomic and transcriptomic annotation were performed employing the version 47 of GENCODE’s human GTF and FASTA files (https://www.gencodegenes.org/human/release_47.html). Reads were aligned with the human genome (GRCh38.p14) by means of STAR (version 2.5.2b) ^48^ and, by using the --outFilterMultimapNmax 1 option, all multimapping reads were discarded. The summarization of paired-end fragments to genes was performed with FeatureCounts (version 1.6.0) ^49^. The process of read alignment with the reference human genome uniquely mapped: i) 82.5% of the ∼1.25 billion input reads, namely ∼84.6 million reads on average per sample. MultiDimensional Scaling (MDS) analyses were performed to study data structure, identify potential outliers, measure the total variability among samples, identifying clusters based on genic expression patterns. MDS analyses were carried out with the regularized log-transformed gene counts (obtained with the DESeq2 *rlog* function) and the *cmdscale* R function; in particular, the aitchison distance was adopted for the MDS. Plots were produced using the ggplot R package (version 3.1.3.1).

For the differential expression analysis, data were prepared via a custom R script. DESeq2 (version 1.26.0) ^50^ was used to perform the normalization of raw sequencing counts, with DESeq2’s mean log ratio method, and the differential expression analysis between the experimental conditions of interest. Before extracting the results of the differential expression analysis, a preliminary gene expression filter was employed and genes, whose sum of normalized counts was less than 10 in half the samples of the dataset were discarded to focus only on highly expressed genes. Genes were considered differentially expressed only when presenting a Benjamini-Hochberg adjusted p-value (false discovery rate, FDR) ≤ 0.05. The functional pathways associated with significant differentially expressed genes were investigated with Ingenuity Pathway Analysis IPA® (Ingenuity Systems, QIAGEN, Redwood City, CA). IPA parameters were kept to their standard values except for species settings (in the species tab, only the Homo sapiens checkbox was considered) and miRNA settings (the box for high confidence predicted miRNAs was checked). The transcript quantifier Kallisto ^51^ was employed to perform the quantification of the *PTBP1* transcripts, by measuring the relative abundance of transcripts as Transcripts Per Million (TPM). The differential alternative splicing analysis was performed with rMATS ^52^ which was used with standard options except for --libType fr-firststrand and --readLength 129. The results obtained were parsed with a custom R script to keep entries with FDR<0.05 and a measured absolute isoform switch equal or greater than 10. Sashimi plots shown in Figure 2D were obtained with rmats2sashimiplot (https://github.com/Xinglab/rmats2sashimiplot).

### ddPCR validation of RNA-seq analysis

RNA-seq data validation was carried out by droplet digital polymerase chain reaction (ddPCR) (Bio-Rad). 200 ng of RNA, extracted from PTBP-siRNA or scramble-siRNA treated hNSCs, were retrotranscribed using the iScript™ Advanced cDNA Synthesis Kit (Bio-Rad Laboratories, cat. #1725038), according to the manufacturer’s instructions. Primer pairs for *PTPB2, GABBR1, FLNA* and *ACTB* were designed using the Isoprimer pipeline ^53^ (PTPB2_for: ACCAATCACAACTTGCCATGA, PTPB2_rev: TGAAGGGTGGCAGAAGGAG; GABBR1_for: CTCACCAGCCCTGTCAAAC; GABBR1_rev: CAGGTCGTCCAGAGTCGAA; FLNA_for: TCGTGGAAGGGGAGAACCAC, FLNA_rev: CTGAATTCGGCTGGCACTCC; ACTB_for: CGCCCTGGACTTCGAGCAAGA, ACTB_rev: CACAGGACTCCATGCCCAGGA). The ddPCR reactions were prepared in a final volume of 22μl, by combining 1 µl of the diluted cDNA template (1:8 for *PTPB2, GABBR1* and *FLNA*, 1:400 for *ACTB*), 11 µl of the 2× QX200™ EvaGreen ddPCR Supermix (Rad Laboratories, cat. # 1864033), primers at a final concentration of 250 nM. and water. Each gene was analyzed at least in triplicate. For each gene, a negative control (No Template Control) was used. Following the manufacturer’s instructions, 20 µl of each ddPCR reaction mix, along with 70 µL of QX200™ Droplet Generation Oil, were loaded into QX200™ Droplet Generator. After droplet generation, samples were transferred into a 96-well plate and amplified using a Bio-Rad T100 Thermal Cycler, with the following profile for all targets: 95 °C for 5 min, 40 cycles of 95 °C for 30 s and 58 °C for 60 s, 4 °C for 5 min, 90 °C for 5 min and hold at 4 °C. The temperature ramp rate was 2 °C/s for all steps. After PCR, the plates were read using a Bio-Rad QX200 Droplet Reader. Absolute quantification was performed using the QuantaSoft version 7.4.1 software (Bio-Rad) and the negative/positive thresholds were set manually. ddPCR reactions were considered positive if characterized by a number of events >10,000, according to the QX200™ reader automatic evaluation. For each sample, results were expressed as the means of transcript copies/μL of PCR replicates, normalized by the means of corresponding ²-actin copies/μL. Normalized counts were analyzed and plotted in GraphPad Prism 10.3.1, using a two-tailed t-test with Welch’s correction. Differences with p < 0.05 were considered statistically significant.

### RT-PCR for alternative splicing analysis

For *PTPB2, GABBR1* and *FLNA* alternative splicing analysis, 200 ng of RNA, extracted from PTBP-siRNA or scramble-siRNA treated hNSCs, were retrotranscribed using the iScript™ Advanced cDNA Synthesis Kit (Bio-Rad Laboratories, cat. #1725038), according to the manufacturer’s instructions. Primer pairs for *PTPB2, GABBR1*, *FLNA* and *ACTB* were designed using the Isoprimer pipeline ^53^ (PTPB2_AS_for: ATGCCTGGAGTCTCAGCTGG, PTPB2_AS_rev: TGAAGGGTGGCAGAAGGAGG; GABBR1_AS_for: TTGTCTCTGGCCATGTGGTGT, GABBR1_AS_rev: AGACTAAAGCCCAGGCCCAG; FLNA_AS_for:ACATACCGCTGCAGCTACCA; FLNA_AS_rev: TCAGCTGTCTCCTTCACCCG; ACTB_for: CGCCCTGGACTTCGAGCAAGA; ACTB_rev: CACAGGACTCCATGCCCAGGA). PCR reactions were carried out in a final volume of 50μl, by combining 1 µl of the cDNA template, primer pairs at a final concentration of 0,5 μM, dNTPs at a final concentration of 0,2 mM, 10U of Taq DNA polymerase (Thermo Scientific, cat.# EP0701) and water. Amplicons were analyzed on 2% of agarose gel (52).

### Mouse brain sections and immunofluorescence

C57BL/6J mice were kindly provided by Professor Giuseppe Calamita, University of Bari Aldo Moro (Authorization from the Italian Ministry of Health No. 158/2024-PR). C57BL/6J mice at postnatal day 15 (P15) were anesthetized by intraperitoneal injection of urethane (1.2 g/kg body weight). Brains were dissected, washed in PBS, and fixed overnight in 4% paraformaldehyde (PFA) in PBS. The following day, brains were washed in PBS, cryoprotected in 30% sucrose in PBS for 24 h, embedded in OCT compound, frozen in liquid nitrogen, and sectioned into 20 µm-thick coronal sections using Superfrost glass slides. Sections were stored at −80°C until use.

Immunofluorescence staining was performed as follows. Sections were rinsed in PBS, fixed for 5 min in 4% PFA, blocked in 3% BSA in PBS for 30 min, and incubated for 48 h at 4°C with the following primary antibodies diluted in 3% BSA and 0.3% Triton X-100 in PBS: goat anti-PTBP1 monoclonal antibody (Sigma, cat. #SAB2500834; 1:1000), guinea pig anti-GFAP antibody (#AFP-001-GP; 1:1000), and rabbit anti-AQP4 antibody (Santa Cruz, sc-9888; 1:500).

After primary antibody incubation, sections were extensively washed in PBS containing 3% BSA and incubated for 24 h at 4°C with appropriate secondary antibodies and Phalloidin Alexa Fluor Plus 750 (Thermo Fisher Scientific, A30105; 1:1000).

Sections were imaged using a Leica Stellaris 8 confocal microscope and images were processed and analyzed using LAS X software.

### Antibodies and working dilution

SOX2 (L1D6A2) Mouse mAb (Cell Signaling, cat #4900) 1:500. Nestin (10C2) Mouse mAb #33475 (Cell Signaling, cat #33475) 1:1000. Anti-mCherry antibody [EPR20579] Rabbit mAb (Abcam, cat #ab213511) 1:500. PTBP1 Goat mAb (Sigma, cat. #SAB2500834) 1:1000. PTBP2 Rabbit mAb (Invitrogen, cat. # PA5-78547) 1:1000. Anti-Ki67 Rabbit mAb (abcam, cat. # ab16667) 1:500. Anti-Flotillin 2 Rabbit Polyclonal Antibody (Novus Biologicals, cat. #NBP130881) 1:1000. CD40/TNFRSF5 Antibody Rabbit Polyclonal Antibody (Novus Biologicals, cat. # NB10056127) 1:1000. Guinea pig anti-GFAP antibody (#AFP-001-GP; 1:1000). Rabbit anti-AQP4 antibody (Santa Cruz, sc-9888; 1:500).

Donkey anti-Rabbit IgG (H+L) Highly Cross-Adsorbed Secondary Antibody, Alexa Fluor™ Plus 488 (Invitrogen, Cat # A32790TR) 1:1000. Donkey anti-Mouse IgG (H+L) Highly Cross-Adsorbed Secondary Antibody, Alexa Fluor™ Plus 488 (Invitrogen, Cat # A32766TR) 1:1000. Donkey anti-Goat IgG (H+L) Highly Cross-Adsorbed Secondary Antibody, Alexa Fluor™ Plus 488 (Invitrogen, Cat # A32814TR) 1:1000. Donkey anti-Rabbit IgG (H+L) Highly Cross-Adsorbed Secondary Antibody, Alexa Fluor™ Plus 555 (Invitrogen, Cat # A32794) 1:1000. Donkey anti-Mouse IgG (H+L) Highly Cross-Adsorbed Secondary Antibody, Alexa Fluor™ Plus 555 (Invitrogen, Cat # A32773) 1:1000. Donkey anti-Goat IgG (H+L) Highly Cross-Adsorbed Secondary Antibody, Alexa Fluor™ Plus 555 (Invitrogen, Cat # A32816) 1:1000. Donkey anti-Rabbit IgG (H+L) Highly Cross-Adsorbed Secondary Antibody, Alexa Fluor™ Plus 647 (Invitrogen, Cat # A32795TR) 1:1000. Donkey anti-Mouse IgG (H+L) Highly Cross-Adsorbed Secondary Antibody, Alexa Fluor™ Plus 647 (Invitrogen, Cat # A32787TR) 1:1000. Donkey anti-Goat IgG (H+L) Highly Cross-Adsorbed Secondary Antibody, Alexa Fluor™ Plus 647 (Invitrogen, Cat # A32849TR) 1:1000. Donkey anti-Guinea Pig IgG, Alexa Fluor™ 647 (Millipore, Cat # AP193SA6), Rabbit anti Goat IgG (H/L): HRP Conjugate (Bio-Rad, cat. #1721034) 1:5000. Goat anti Rabbit IgG (H/L): HRP Conjugate (Bio-Rad, cat. #1706515) 1:5000. Goat anti Mouse IgG (H/L): HRP Conjugate (Bio-Rad, cat. #1706516.) 1:5000.

### Software for illustration generation

The graphical abstract, Figure 3G, and the cartoon shown in Figure 9 were created using BioRender. Agreement number: PP29DPPZIP.

### Experimental design and statistical analysis

All data represent at least three independent experiments for each experimental group (see figure legends for details). Statistical analyses were performed using GraphPad Prism 9 software (GraphPad Software). Data are presented as mean ± SEM. Statistical significance was assessed using an unpaired Student’s t-test or two-way ANOVA followed by Tukey’s multiple-comparisons test, as appropriate. A p value < 0.05 was considered statistically significant.

## Results

### PTBP1 regulates hNSC fate by controlling mitochondria, lipid droplets, and PTBP2 expression

The functional role of PTBP1 was investigated through siRNA-mediated PTBP1 knockdown (KD) using a pooled mix targeting all known PTBP1 isoforms. All experiments were conducted by maintaining hNSCs in a hypoxic incubator, using a medium specifically formulated to preserve hNSC stemness ^44^.

Western blot analysis confirmed efficient PTBP1 KD in monolayer cultures at 7 days in vitro (DIV) (Figure 1A). Cell counts at 7 DIV revealed that PTBP1 KD significantly reduced the number of cells compared to control siRNA treatment (Figure 1B).

Knockdown efficiency was also confirmed in neurospheres at 7 DIV (Figure 1C). PTBP1 KD led to a marked reduction in the number of floating neurospheres, total cell count, and neurosphere volume, while increasing the number of adherent cells compared to the control siRNA condition (Figure 1D). Neurospheres at 7 DIV, treated with control or PTBP1 siRNA for the entire 7-day period, were mechanically dissociated and seeded as a single-cell suspension onto Cultrex-coated coverslips. Cells were analyzed for Ki67 expression the following day, maintaining siRNA in the culture medium. Immunofluorescence analysis showed that PTBP1 knockdown decreased the percentage of Ki67⁺ cells (Figure 1E).

Neurospheres at 7DIV treated with control or PTBP1 siRNA were mechanically dissociated and analyzed at the single-cell level for mitochondrial membrane potential using the ratiometric dye JC-1, and by live-cell, label-free holotomographic imaging to quantify the number, dry mass, and morphology of mitochondria and lipid droplets (LDs) under physiological hypoxia. This analysis revealed that PTBP1 KD increased mitochondrial membrane potential (Figure 1F), as well as the number and mitochondrial dry mass per cell, while reducing mitochondrial length and area (Figure 1G). Furthermore, PTBP1 KD decreased both the number and dry mass of LDs per cell (Figure 1H). Finally, PTBP1 KD strongly upregulated PTBP2 expression (Figure 1I).

Considering the key role of PTBP2 in neuronal differentiation of NSCs ^54^ and the emerging involvement of mitochondria^55^, and LDs^56^ in NSC fate commitment toward neurons, these data collectively demonstrate that PTBP1 regulates hNSC fate at multiple levels, from cell proliferation to cellular metabolism, maintaining cells to a stem-related state.

**Figure 1.**
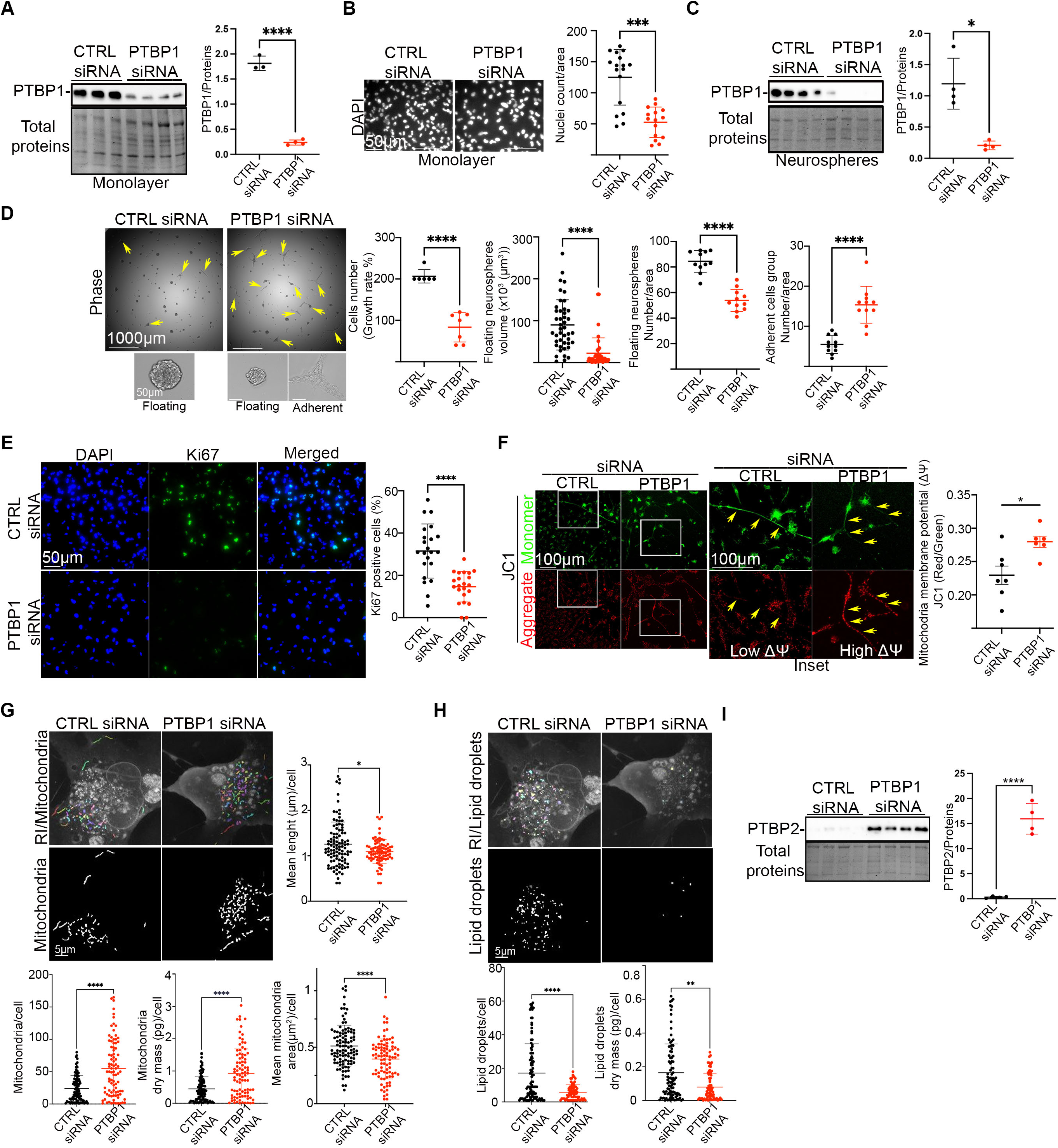
PTBP1 knockdown alters hNSC fate by regulating mitochondria, lipid droplets, and PTBP2 expression. **A.** Western blot analysis of PTBP1 expression in control (CTRL) and PTBP1 siRNA–treated hNSCs cultured as monolayers for 7 days in vitro (DIV) shows efficient PTBP1 knockdown (KD). Each lane represents an independent experiment (n=3-4 for each group). **B.** Quantification of DAPI-stained nuclei/area at 7 DIV in CTRL and PTBP1 siRNA conditions shows that PTBP1 KD significantly reduces cell number. Representative images. Each point represents the cell number per random field (n = 3 for each group). **C.** Western blot analysis of PTBP1 expression in CTRL and PTBP1 siRNA–treated hNSC neurospheres at 7 DIV confirms efficient KD. Each lane represents an independent experiment (n=4 for each group). **D.** Neurosphere assays show that PTBP1 KD markedly decreases total cell content, the number of floating neurospheres, and neurosphere volume, while increasing the number of adherent cell clusters. Each point represents the total cell count of mechanically dissociated neurospheres, the number of neurospheres, neurosphere volume, or the number of adherent clusters per random field (n = 3–5 for each group. **E.** Immunofluorescence analysis of Ki67 in hNSCs treated with control or PTBP1 siRNA. PTBP1 KD reduces the percentage of Ki67⁺ cells. Each point represents the percentage of Ki67⁺ cells per random field (n = 3–5 for each group). **F.** Analysis of mitochondrial membrane potential using JC-1. Red fluorescence (JC-1 aggregates) indicates high membrane potential, while green fluorescence (JC-1 monomers) indicates low membrane potential. The red/green fluorescence ratio measures mitochondrial membrane potential. PTBP1 KD increases mitochondrial membrane potential. Each point represents the average red/gree ratio value per random field (n = 3 for each group). Arrows indicate low and high ²q, mitochondria in CTRL and PTBP1 siRNA treated hNSCs, respectively. **G.** Label-free holotomographic morphometric analysis of mitochondria in hNSC neurospheres at 7 days in vitro (DIV) following control or PTBP1 siRNA treatment. PTBP1 KD significantly increases mitochondrial number and dry mass per cell, while reducing mitochondrial length and area. Each point represents a single cell. Total cells analyzed = 33–125 per group (n = 3 for each group). **H.** Label-free holotomographic morphometric analysis of lipid droplets (LDs) in hNSC neurospheres at 7 DIV (CTRL siRNA vs. PTBP1 siRNA). PTBP1 KD significantly reduces LD number and dry mass per cell. Each point represents a single cell. Total cells analyzed = 33–125 per group (n = 3 for each group). **I.** Western blot analyses of PTBP2 expression in CTRL and PTBP1 siRNA–treated hNSCs. PTBP1 KD strongly upregulates PTBP2 expression. Each lane represents an independent experiment (n=4 for each group).

### PTBP1 controls the expression profiling of hundreds of coding and non-coding genes, regulating stemness and neuronal differentiation

To assess the role of PTBP1 in hNSC fate regulation at the RNA level, we performed bulk RNA sequencing (RNA-seq) on hNSCs treated with control or PTBP1 siRNA. The differentially expression analysis, performed by DESeq2, identified 248 deregulated genes between PTBP1 siRNA treated hNSCs and control siRNA treated cells (FDR<0.05), of which 221 were protein coding genes (102 upregulated and 119 downregulated) and 27 were genes for long non coding RNAs (lncRNAs) (17 upregulated and 10 downregulated) (Figure 2A and Supplementary Table 1). The top upregulated and downregulated genes are reported in Figure 2B. Interestingly, we observed that the knockdown of PTBP1 induced the overexpression of its paralog *PTBP2* and of *GABBR1* and the repression of the expression of *FLNA* (Figure 2C). In particular, we show that the expression of these genes is regulated by PTBP1 through alternative splicing, via repression of cassette exon inclusion in their pre-mRNAs, similar to what has been reported in other cell types ^57–59^ (Figure 2D). We found that PTBP1 knockdown induces the inclusion of 34 nt long exon in *PTBP2* mRNA and of 151 nt long exon in *GABBR1* mRNA, thus preventing nonsense-mediated decay (NMD) of these transcripts, and explaining the strong upregulation found for both genes in PTBP1-siRNA treated hNSCs (Figure 2E and Figure 1I). On the contrary, we observed that PTBP1 knockdown induces the inclusion of a conserved 57 nt long exon in the *FLNA* transcript who demonstrated that this exon produces degradation of the murine *FLNA* transcript by NMD, thus explaining the strong downregulation found for *FLNA* in PTBP1 siRNA treated hNSCs (Figure 2E). Differentially expressed genes then were examined with Ingenuity Pathway Analysis (IPA) to identify significant pathways deregulated in PTBP1-siRNA treated hNSCs; gene modules were considered to be associated with a canonical pathway if the Fisher’s exact test performed by IPA had a p-value < 0.05. As shown in Figure 2F, among the upregulated pathways, we found signaling pathways (e.g., acetylcholine receptor signaling and protein kinase A signaling) and the human embryonic stem cell pluripotency pathway; among the downregulated pathways, we found pathways involved in post-translational protein phosphorylation, in the regulation of insulin-like growth factor (IGF) transport and uptake by IGFBPs and in ion channel transport. Considering the central role of PTBP2 as a master regulator of neuron-specific alternative splicing programs ^60,61^, the pivotal function of GABBR1 in GABAergic neuron–mediated neural circuits ^62^, and the role of FLNA in regulating radial glial and neural progenitor cell fate ^63–65^, the results presented here demonstrate that PTBP1 represses neuronal differentiation and preserves the stemness state of hNSCs by regulating hundreds of genes, including key neurogenesis-associated factors, through both transcriptional and splicing-dependent mechanisms.

**Figure 2.**
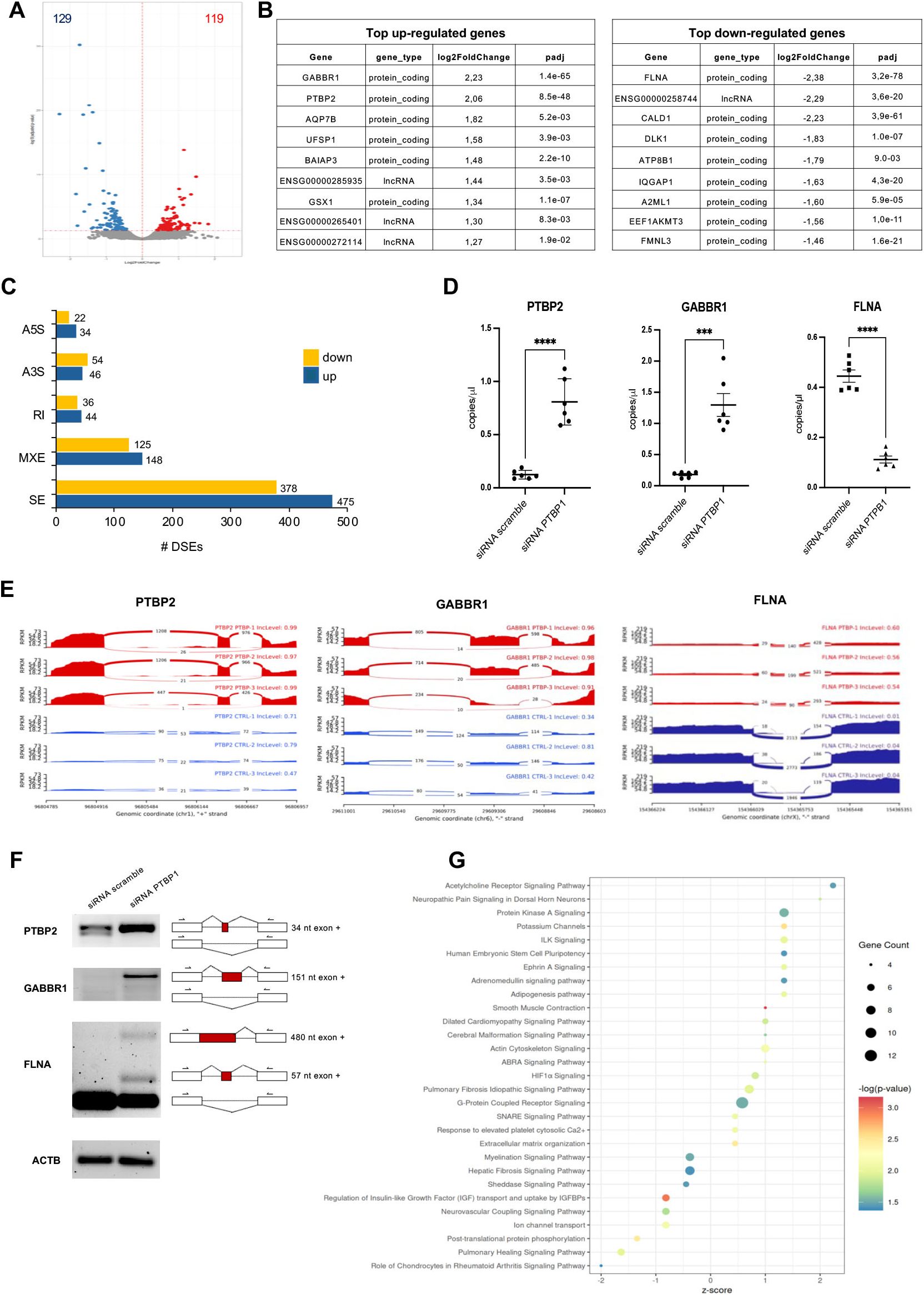
PTBP1 knockdown alters hNSC gene expression profiling. **A**. Volcano plot of differentially expressed genes (DEGs) in PTBP1 siRNA treated hNSCs versus scramble siRNA treated cells. The statistically significant (False Discovery rate, FDR ≤ 0.05) up-regulated and down-regulated genes are shown in red and blue, respectively (n=3 for each group). **B.** Top most significantly deregulated genes, with *GABBR1 and PTBP2* among the top up regulated, and *FLNA* among the most down regulated genes. The log2FoldChange and the Benjamini-Hochberg adjusted p-value (padj) are reported. **C.** Bar plot showing the number of differentially spliced events (DSE) identified in five types of alternative splicing events. SE: skipped exon; A5S: alternative 5’ splice site; A3S: alternative 3’ splice site; MXE: mutually exclusive exons; RI: retained intron. **D.** RT-qPCR validation, by ddPCR, of *GABBR1, PTBP2* and *FLNA* expression in PTBP1 siRNA treated hNSCs versus scramble siRNA treated hNSCs. Data are represented as transcript copies/μL of reaction mix, normalized with respect to *²-Actin* transcript copies/μL. Two-tailed t-test with Welch’s correction was used. ***p<0.001, ****p<0.0001. (n=3 for each group). **E.** Sashimi plots of PTBP1 responsive exons in *GABBR1, PTBP2* and *FLNA* transcripts in PTBP1 siRNA treated (in red) and scramble siRNA treated (in orange) hNSCs. RPKM is plotted on the y-axis. Each x-axis represents a replicate (n=3 for each group). **F.** RT-PCR validation of alternatively spliced exons of *GABBR1, PTBP2* and *FLNA* transcripts in PTBP1 siRNA treated and scramble siRNA treated hNSCs, reported in the Sashimi plots of panel D (n=3 for each group). **G.** Functional pathways deregulated in PTPB1 siRNA treated hNSCs investigated with Ingenuity Pathway Analysis (IPA). Z-scores ≥0 represent predictions of activation, Z-scores ≤ 0 represent predictions of inhibitions (n=3) for each group (65).

### The overexpression of PTBP1 *isoform a* alters the endogenous PTBP1 isoform pattern and changes hNSC fate

RNA-seq data were analyzed to characterize the expression profile of *PTBP1* transcripts in untreated hNSCs. We found that hNSCs predominantly express three *PTBP1* transcripts: ENST00000587191, ENST00000356948 and ENST00000349038, although additional *PTBP1* mRNAs, expressed at a very low level, were also detected (Figure 3A and Supplementary Table 1). The first transcript (ENST00000587191) has an average Transcripts Per Million (TPM) across samples of 83.88 and should encode a never characterized protein of 81 amino acids (aa). The other two most abundant transcripts differ in the inclusion or exclusion of exon 9. The ENST00000356948 transcript has an average TPM across samples of 33.37. It includes exon 9 and encodes a protein of 557 aa, also known as PTBP1 *isoform a* (NP_002810.1). The ENST00000349038 transcript has a measured average TPM of 19.52 and encodes a protein of 531 aa, known as PTBP1 *isoform c* (NP_114368.1) (Figure 3A, red arrows).

To assess PTBP1 isoform expression at the protein level, high-resolution Western blotting was performed. Consistent with the RNA-seq results, hNSCs exhibited two main protein bands corresponding to the *isoform a* and *isoform c* and the PTBP1 *isoform a*, was found to be the predominant isoform at protein level (Figure 3B).

To determine how hNSCs respond to the perturbation of the physiological PTBP1 isoform ratio, the coding sequence of PTBP1 *isoform a* was cloned into a lentiviral (LV) vector to express an mCherry-tagged protein. hNSCs were transduced with PTBP1-mCherry LV, with mCherry LV used as a control. Western blot analysis revealed that the ectopic expression of PTBP1 *isoform a*-mCherry induced a marked downregulation of both endogenous PTBP1 *isoform a* and *c*, compared with the control virus (Figure 3C). In addition, a new band induced by PTBP1 *isoform a* appeared, compatible with isoforms encoded by ENST00000635647 (588 aa) and ENST00000893849 (606 aa) transcripts (Figure 3C, indicated by “X”).

hNSCs expressing mCherry or PTBP1 *isoform a*–mCherry were analyzed for Ki67⁺ cells and Nestin expression. Both markers were significantly reduced in PTBP1 *isoform a*–overexpressing cells compared with control cells. Mitochondrial membrane potential, assessed using JC-1, showed a strong increase, upon PTBP1 *isoform a* overexpression. Label-free holotomographic analysis of mitochondrial and lipid droplet morphometry further showed that both mitochondrial and lipid droplet content were reduced in the PTBP1 *isoform a*-mCherry LV group compared with the mCherry LV controls.

Taken together, these findings demonstrate that overexpression of PTBP1 *isoform a* profoundly alters the endogenous PTBP1 isoform landscape, reducing proliferation and stemness markers while promoting mitochondria and lipid droplets alteration characteristic of early differentiation ^43,73^—despite culture in hypoxia and in a medium specifically formulated to preserve hNSC stemness ^44^. These results highlight that hNSC stemness is compromised when the physiological PTBP1 isoform pattern is perturbed.

**Figure 3.**
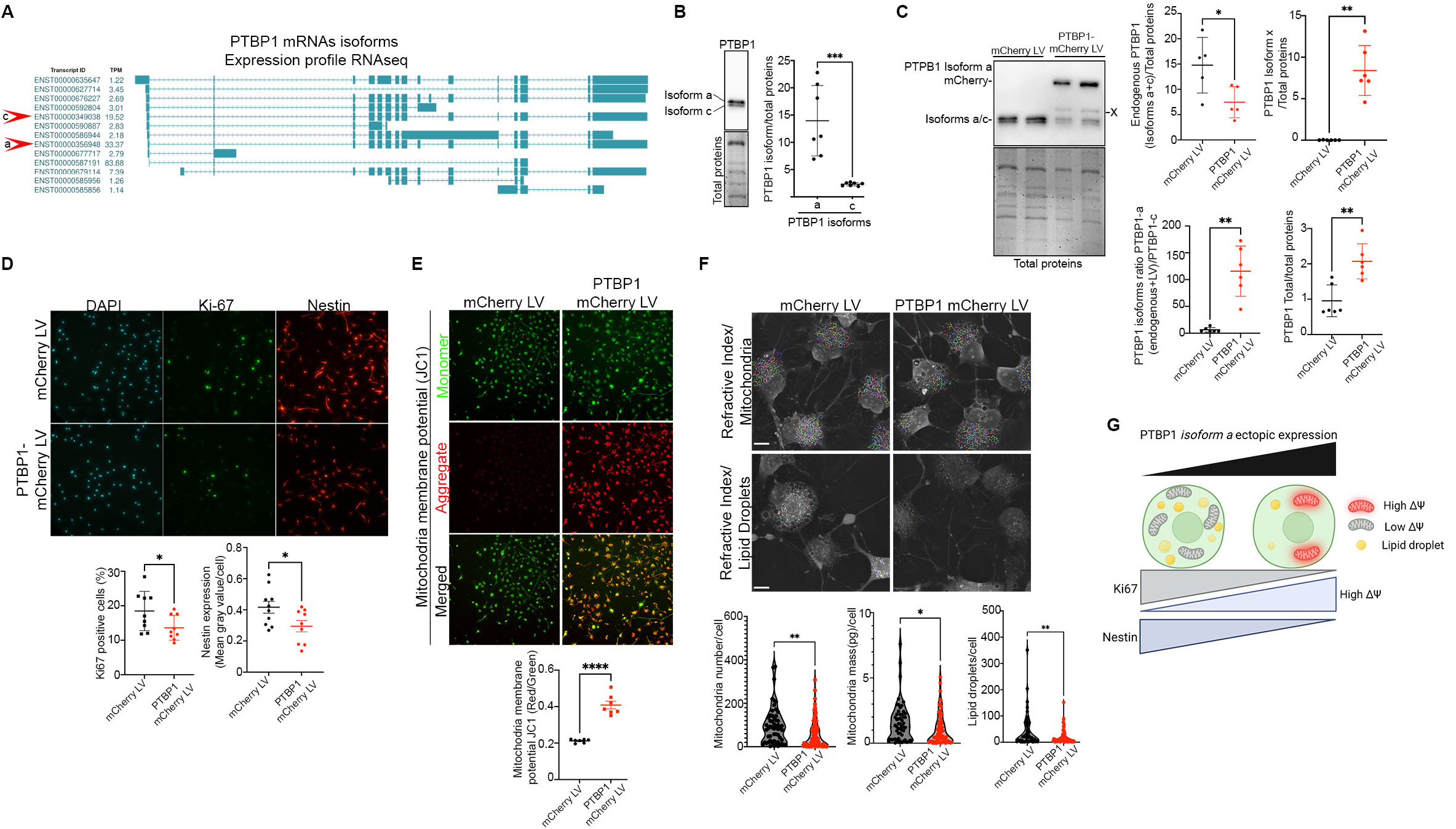
The overexpression of PTBP1 *isoform a* alters the endogenous PTBP1 isoform pattern and changes hNSC fate. **A.** Expression profile of *PTBP1* transcripts derived from RNA-seq of non-treated hNSCs. Transcript quantitation was performed with Kallisto tool ^51^ using the version 47 of the human GENCODE genomic annotation. The Ensembl ID and the measured average Transcripts Per Million (TPM) of each transcript across samples are reported. Only transcripts with TPM >1 are indicated. Major isoforms (a and c) are indicated by red arrowhead. **B.** Western blot analysis showing two major endogenous PTBP1 protein bands corresponding to the 557 aa and 531 aa isoforms (*isoforms a* and *c,* respectively). Representative Western blotting. Each point represents an independent sample (n=7). **C.** Western blotting of hNSCs transduced with control mCherry LV or PTBP1 *isoform a*–mCherry LV. Overexpression of PTBP1 *isoform a* drastically alters the endogenous PTBP1 isoform pattern and disrupts the physiological isoform ratio. Representative Western blotting. Each point represents an independent experiment (n=5-6 for each group). **D.** Immunofluorescence for Ki67 and Nestin in hNSCs transduced with mCherry or PTBP1 *isoform a*–mCherry. PTBP1 *isoform a* overexpression reduces the percentage of Ki67⁺ cells and decreases Nestin expression. Each point represents the percentage of Ki67⁺ cells per random field or nestin mean fluorescence intensity per random field (n = 3 for each group). **E.** Mitochondrial membrane potential analysis using the JC-1 dye. Overexpression of PTBP1 *isoform a*–mCherry strongly increases mitochondrial membrane potential. Each point represents the average value per random field (n = 3 for each group). **F.** Label-free holotomographic morphometric analysis of mitochondria and lipid droplets (LDs) in hNSCs transduced with control mCherry LV or PTBP1 *isoform a*–mCherry LV. PTBP1 overexpression alters mitochondrial and LD abundance and morphology. Each point represents a single cell. Total cells analyzed = 90–95 per group (n = 3 for each group). **G.** Proposed model. PTBP1 isoform a overexpression alters hNSC stemness and metabolic state, reducing cell proliferation, Nestin expression, lipid droplet abundance, and mitochondrial content, while increasing mitochondrial membrane potential.

### PTBP1 Localizes to the Nucleus, Cytoplasm, TNTs, Extracellular Vesicles, and Migrasomes in hNSCs

To investigate the subcellular localization of PTBP1 in hNSCs, we performed immunofluorescence and τ-STED super-resolution confocal microscopy for PTBP1, F-actin, and nestin. Our analysis revealed that hNSCs are highly interconnected by TNTs and that PTBP1 localizes not only to the nucleus but also to the cytoplasm and along TNTs (Figure 4A–B). Super-resolution τ-STED microscopy showed PTBP1 within the cytoplasm and TNTs as discrete, spot-like signals with areas of approximately 1 × 10³ nm² and integer multiples thereof, indicative of highly ordered structures. The majority of cytosolic PTBP1 clusters ranged from 1 to 4 × 10³ nm² (Figure 4C). To study PTBP1 dynamics in living hNSCs, neurospheres expressing PTBP1 (*isoform a*)–mCherry were imaged using live-cell time-lapse holotomography combined with fluorescence under physiological hypoxic conditions. Time-lapse imaging of migrating LV-transduced neurospheres revealed three distinct and highly dynamic localization patterns of PTBP1-mCherry: nuclear, cytosolic, and dual localization (Figure 5A, insets a and b). Within a few hours after the onset of migration, we observed PTBP1-mCherry “surfing” along TNTs, floating within extracellular vesicles (EVs) (Figure 5B. Supplementary Video 1-2), and being released in vesicles generated by retraction structures resembling migrasomes^36^ (Figure 5C. Supplementary Video 3). Notably, PTBP1-mCherry–containing vesicles were also actively captured by migrating hNSCs (Figure 5D, Inset. Supplementary Video 4).

Together, these findings demonstrate that PTBP1 is not confined to the nucleus but also localizes to the cytoplasm and traffics through TNTs, EVs, and migrasomes, supporting the hypothesis of intercellular transfer among hNSCs.

**Figure 4:**
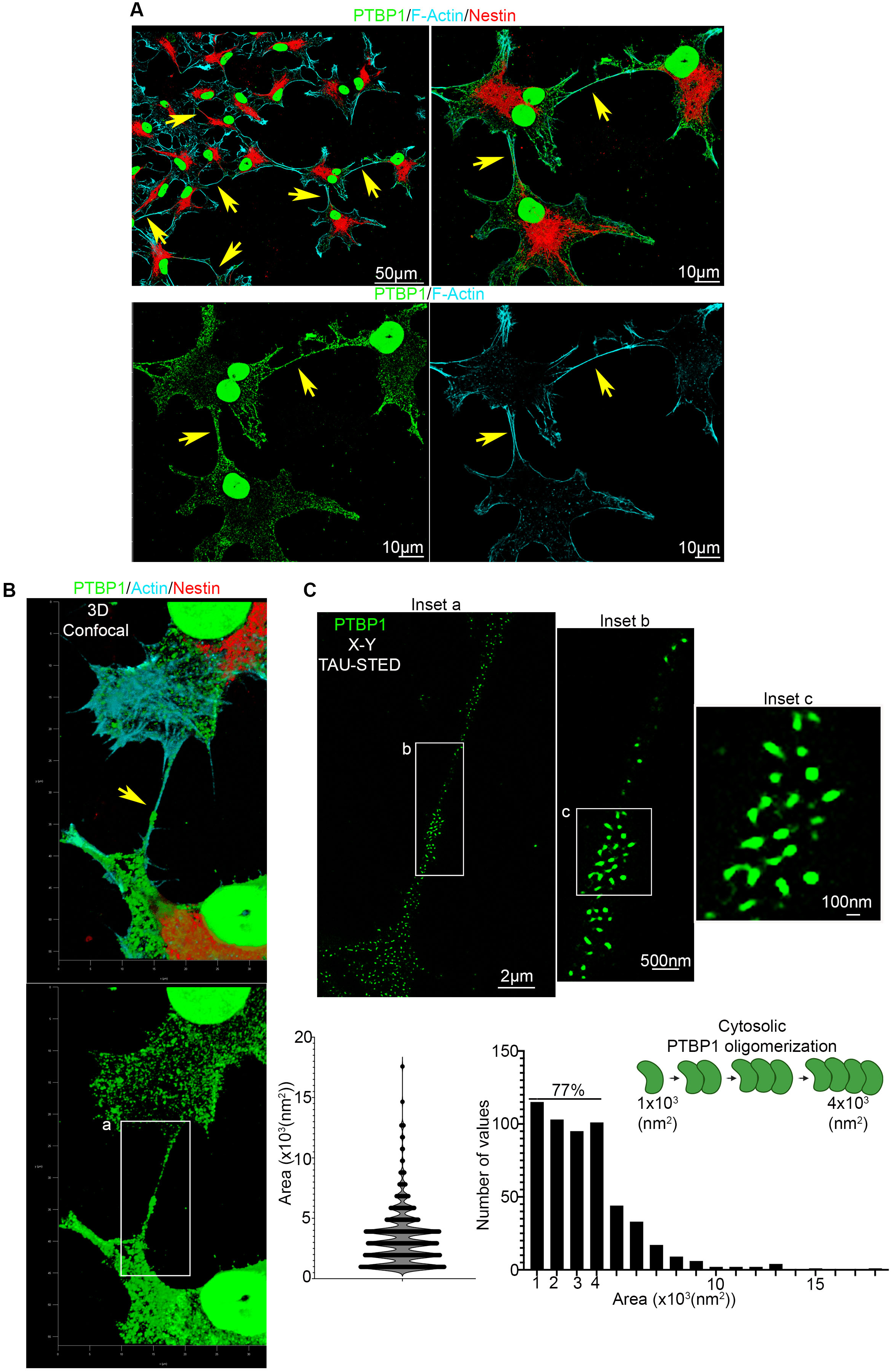
PTBP1 localizes to the nucleus, cytoplasm, and TNTs as highly ordered structures. **A–B**. Confocal microscopy of migrating neurospheres (A) and 2D cultures of hNSCs (B) shows that PTBP1 localizes to the nucleus, the cytoplasm, and within F-actin/Nestin–positive TNTs connecting hNSCs. Representative images (n=6 independent experiments). Arrows indicate PTBP1-containing TNTs. **C.** Super-resolution τ-STED microscopy revealed PTBP1 localization within the cytoplasm and TNTs as discrete, spot-like signals with areas corresponding to multiples of ∼1 × 10³ nm², indicative of highly ordered structures. Seventy-seven percent of the spots were multimers of a single unit, comprising up to four units. Each point represents one PTBP1 cluster (N = 535 clusters, n = 3 independent experiments). Arrows indicate PTBP1-containing TNTs.

**Figure 5:**
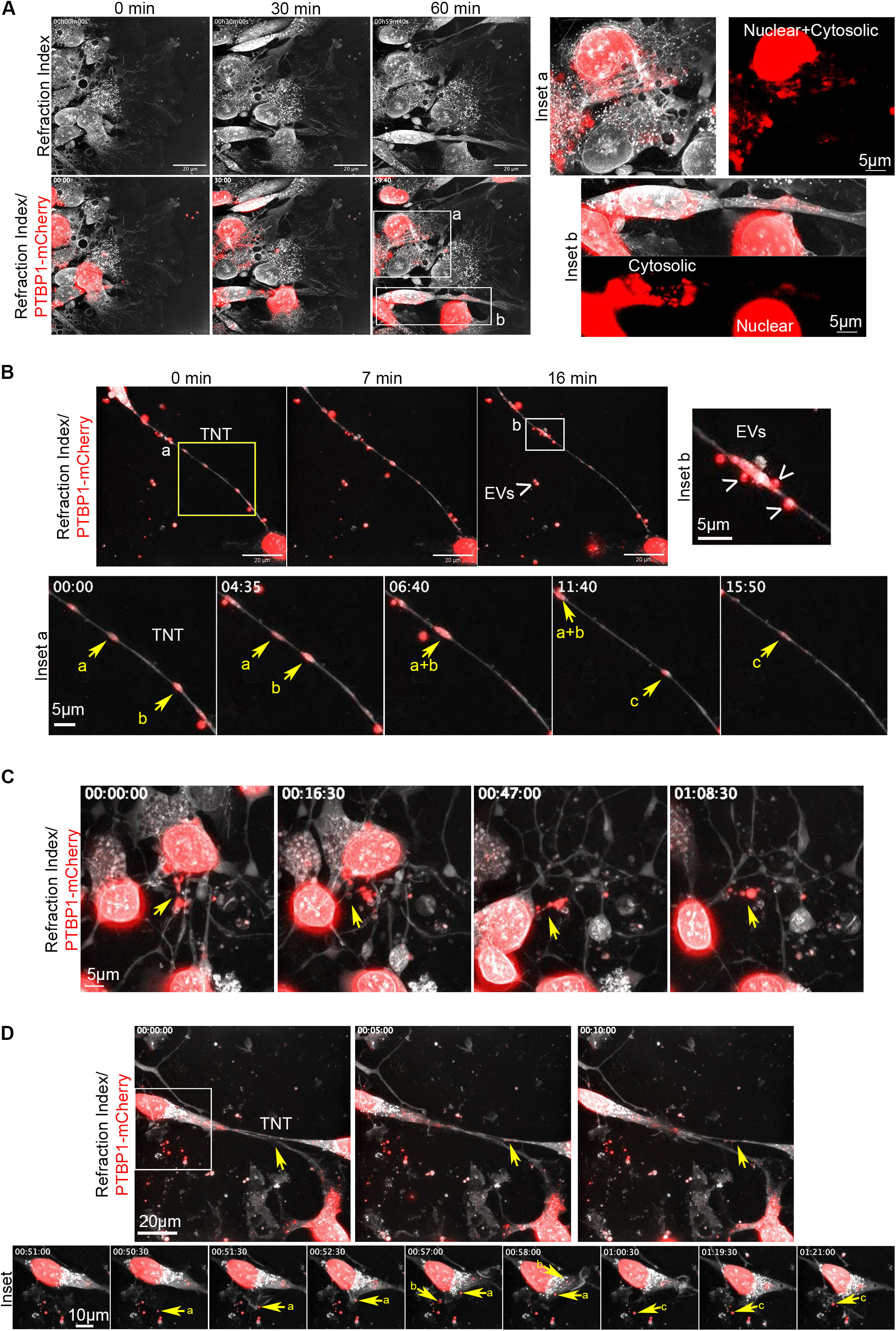
PTBP1 Moves within TNTs and Is Released into EVs and Migrasomes. **(A–D)** Holotomographic and fluorescence time-lapse imaging of LV-transduced neurospheres reveals the highly dynamic behavior of PTBP1-mCherry during neurosphere migration, showing both nuclear and cytosolic localization (A). Within a few hours, PTBP1-mCherry localizes within TNTs EVs, trafficking through these structures and moving between cells (yellow arrows) (B) Supplementary Video 1-2. During cell migration, we observed the formation of a large number of vesicles generated by retraction structures resembling migrasomes (C). Supplementary Video 3. An hNSC actively captures EVs released by another cell (D). Supplementary Video 4. n = 5 independent experiments.

### PTBP1 Is Efficiently Transferred between hNSCs via TNTs

To evaluate whether PTBP1 can be transferred between hNSCs, we performed direct-contact co-culture experiments using hNSCs expressing the dominant PTBP1 isoform-a mCherry as donor cells and MitoTracker-labeled hNSCs as acceptor cells. Live-cell holotomography/fluorescence imaging under physiological hypoxic conditions revealed the movement of PTBP1–mCherry from donor to acceptor hNSCs through TNTs (Figure 6A. Supplementary Video 5-6). Confocal microscopy of co-cultures fixed 24 h later confirmed the presence of PTBP1–mCherry within F-actin–positive TNTs connecting donor and acceptor cells, its transfer into the cytoplasm of acceptor cells, and its localization within acceptor-cell nuclei identified by Sox2 staining (Figure 6B).

To quantify PTBP1–mCherry intercellular transfer, PTBP1–mCherry–expressing hNSCs were co-cultured in direct contact with MitoTracker-labeled or unlabeled recipient hNSCs at a 1:1 cell ratio. In parallel, non-contact co-cultures were established using Transwells with a 400-nm pore size to assess EV-mediated transfer, maintaining the same cell ratio. Experiments were performed for 24 h or 72 h to evaluate temporal dynamics. Recipient cells cultured alone served as negative controls. All cells were stained for Sox2, and PTBP1–mCherry was detected by anti-mCherry immunofluorescence. Samples were analyzed by quantitative confocal microscopy.

PTBP1–mCherry was detected in the cytoplasm of acceptor cells under both direct-contact and non-contact conditions; however, nuclear transfer was time dependent. Quantification of PTBP1–mCherry within nuclei of acceptor hNSCs showed that direct-contact co-culture supported rapid nuclear transfer within 24 h (Figure 6C), whereas non-contact co-culture resulted in detectable nuclear localization only after 72 h (Figure 6D).

To determine whether intercellular transfer mediated by direct contact depends on PTBP1 levels in recipient cells, PTBP1–mCherry–expressing hNSCs were co-cultured with acceptor hNSCs treated with either control siRNA (CTRL-siRNA) or PTBP1-targeting siRNA (PTBP1-siRNA). Quantitative analysis revealed that nuclear transfer of PTBP1–mCherry was approximately twofold higher in PTBP1KD recipient cells (Figure 7A).

We next investigated whether the extensive nuclear transfer of PTBP1 observed in PTBP1 KD cells was sufficient to rescue the reduction in proliferation (Ki67+) induced by PTBP1 depletion (Figure 1E). Given the detrimental effects of PTBP1–mCherry expression on hNSC fate and endogenous PTBP1 isoforms (Figure 3), this experiment was performed using direct-contact co-cultures between normal hNSCs (donors) and PTBP1 KD hNSCs (recipients). A 400-nm pore size Transwell-based non-contact co-culture was used as a control for direct contact to evaluate potential EVs-mediated transfer. Because PTBP1 knockdown upregulates PTBP2 expression (see Figure 1), PTBP2 staining was used to identify PTBP1 KD acceptor cells. Co-cultures were fixed after 72 h and analyzed by quantitative confocal microscopy for Ki67 in PTBP1 KD acceptor cells (PTBP2^+^).

Quantitative analysis revealed no significant difference in Ki67 positivity between PTBP1 KD cells cultured alone and those in direct-contact co-culture. In contrast, non-contact co-culture resulted in a significant increase in Ki67 positivity in acceptor cells respect PTBP1 KD alone (Figure 7B). High-magnification imaging of direct-contact co-cultures revealed high levels of PTBP2 in the cytoplasm of PTBP1 KD cells and, notably, within TNTs connecting PTBP1 KD cells with normal hNSCs, as well as within the cytoplasm and nucleus of normal hNSCs (Figure 7C-D). These observations indicate that while normal hNSCs transfer PTBP1 to PTBP1 KD cells through direct contact, PTBP1 KD cells reciprocally transfer PTBP2 to normal hNSCs, potentially counterbalancing the proliferative rescue expected from PTBP1 transfer.

This mechanism may explain the lack of Ki67 recovery observed in PTBP1 KD cells under direct-contact conditions and the partial recovery observed in non-contact co-cultures, likely mediated by EV-associated PTBP1 transfer.

Together, these data demonstrate that PTBP1 transfer between hNSCs is mediated by direct cell–cell contact and suggested also an EVs-mediate trafficking.

**Figure 6.**
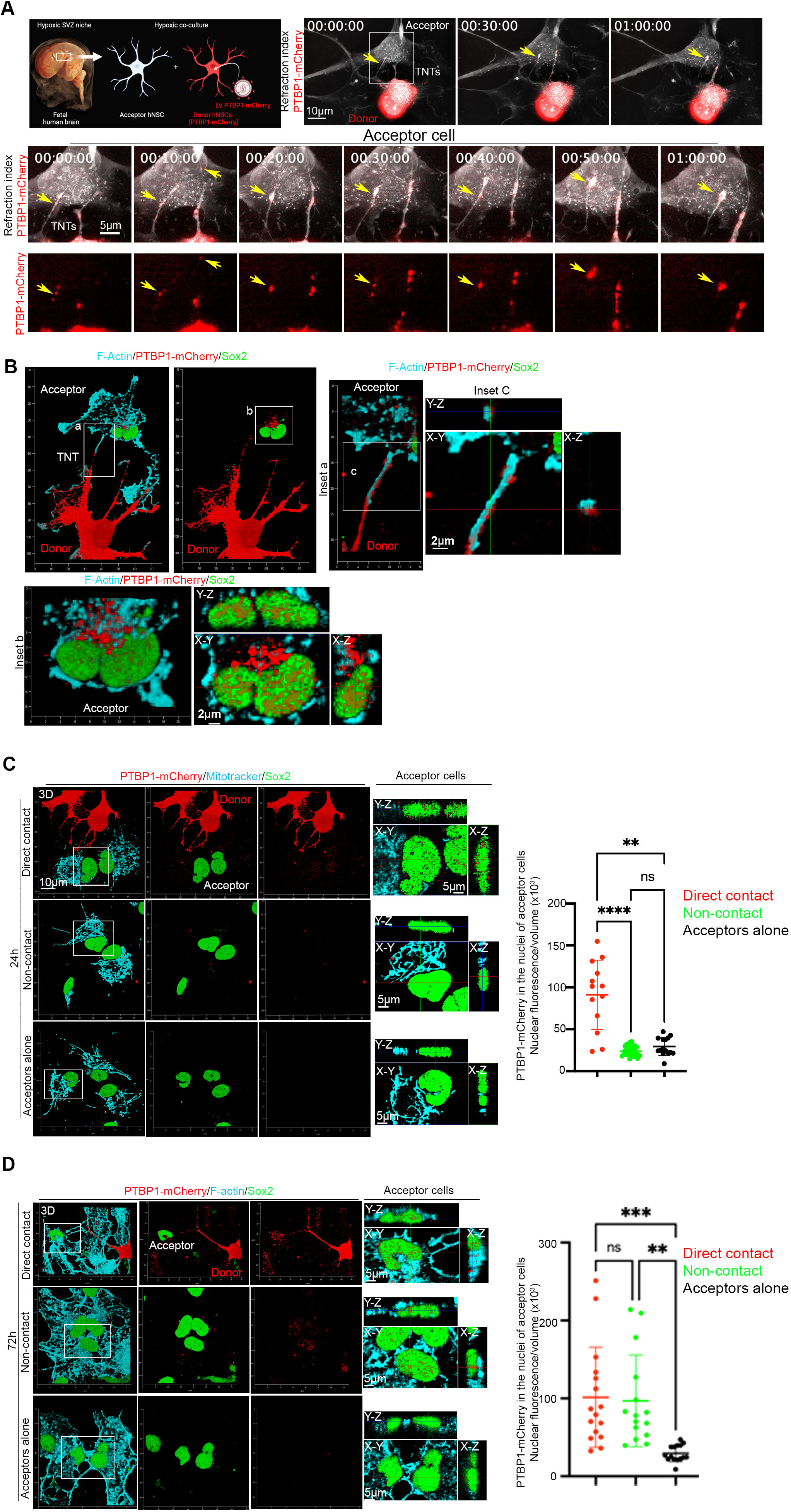
PTBP1 is rapidly transferred to the nuclei of recipient hNSCs via TNTs. **A.** Representative holotomographic/fluorescence time-lapse imaging of a co-culture of PTBP1–mCherry–expressing hNSCs with unstained recipient cells. TNTs connecting donor and recipient cells are visible, and PTBP1–mCherry is observed trafficking from the donor to the recipient cell through a TNT (n=4) (Supplementary Video 5-6). **B.** Representative confocal microscopy of the co-culture shown in (A) confirms the presence of F-actin–positive TNTs connecting donor and recipient cells, through which PTBP1–mCherry is transferred. Analysis of the recipient cell nucleus (Sox2-positive) clearly shows PTBP1–mCherry localization within the nucleus (n=6). **C–D.** Quantitative analysis of nuclear PTBP1–mCherry signal in Sox2-positive, MitoTracker-labeled recipient cells under direct-contact and non-contact co-culture conditions. Direct contact results in higher levels of PTBP1 transfer into recipient cell nuclei within 24 h (C), whereas non-contact co-culture mediates nuclear transfer at 72 h (D). Each point represents the average nuclear PTBP1–mCherry fluorescence normalized to cell volume in each random field (n = 3 for each group).

**Figure 7:**
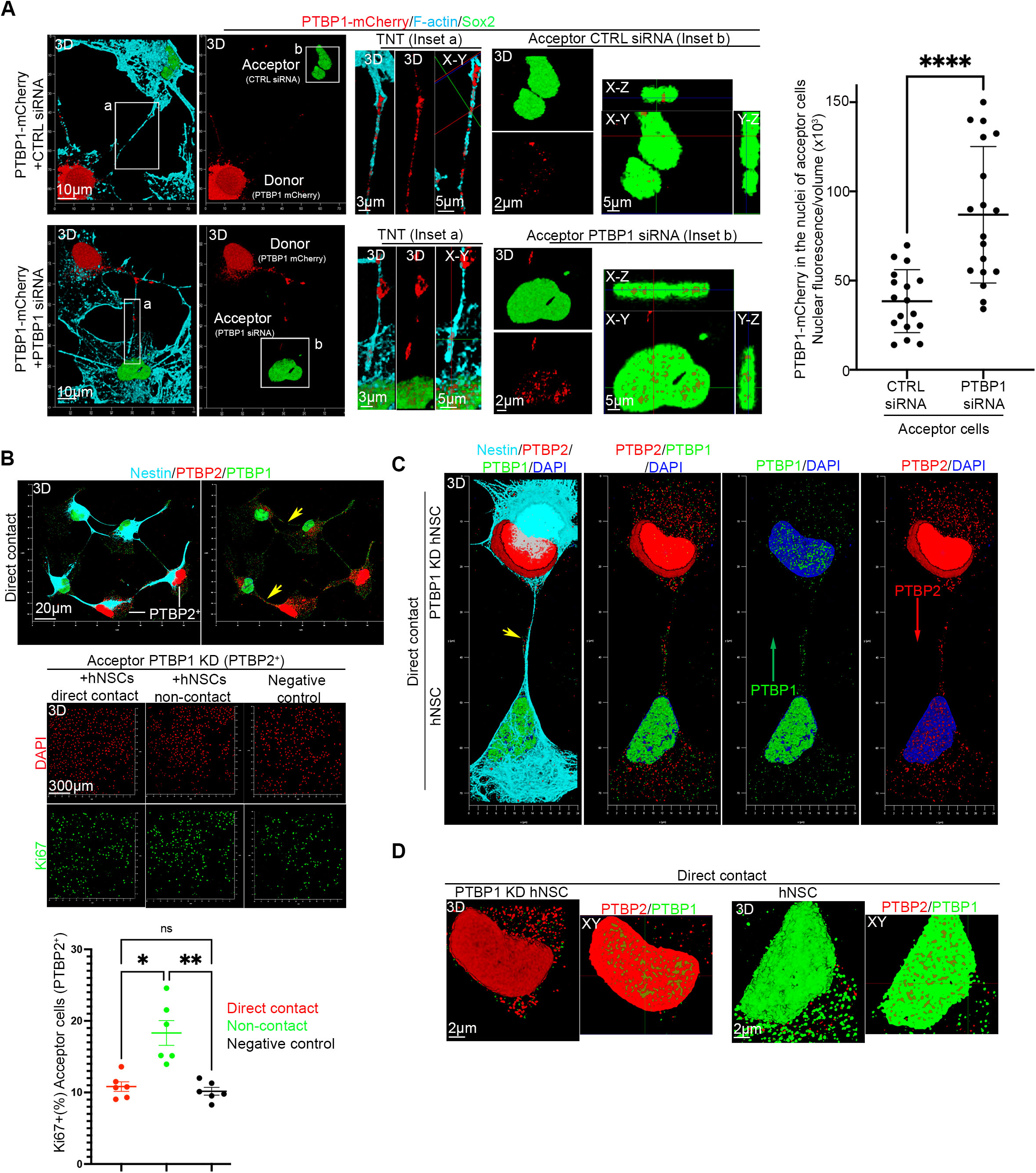
Intercellular transfer mediated by TNTs depends on PTBP1 levels in recipient cells, while reciprocal PTBP1/PTBP2 exchange may prevent proliferation recovery. **A.** Intercellular transfer mediated by direct contact depends on PTBP1 levels in recipient cells. Representative images of direct-contact cocultures between PTBP1–mCherry donor cells and CTRL or PTBP1-siRNA recipient cells. PTBP1–mCherry is visible in TNTs connecting donor and recipient cells and inside nucleus (Sox-2) of recipient cells. Quantitative analysis shows that nuclear transfer of PTBP1–mCherry is approximately twofold higher in PTBP1-depleted acceptor cells. Each point represents the average nuclear PTBP1–mCherry fluorescence normalized to cell volume in each random field (n = 3 for each group). **B.** Analysis of Ki67 positivity in acceptor cells under direct-contact, non-contact, and negative control conditions. Direct contact does not promote recovery of Ki67 positivity in PTBP1-KD acceptor cells (PTBP2-positive cells), whereas non-contact coculture leads to an increase in Ki67 positivity. Each point represents the percentage of Ki67-positive cells per random field (n = 3 for each group). **C.** High-magnification and super-resolution confocal microscopy of direct-contact cocultures between normal hNSCs (PTBP1-positive/PTBP2-negative) and PTBP1-KD cells (PTBP2-positive/PTBP1-negative). TNTs connecting the two cell types show potential reciprocal exchange of PTBP1 and PTBP2. PTBP1 (green), PTBP2 (red). **D.** High-magnification images with optical sectioning of nuclei from normal and PTBP1-KD cells. PTBP1 (green) is detected in the nuclei of PTBP1-KD cells (PTBP2-positive, red), while PTBP2 (red) is detected in the nuclei of normal cells (PTBP1-positive, green).

### Large and micro EVs Contain Specific, Uncharacterized PTBP1 Isoforms and EV Transfer Regulates Cell Proliferation

Large and micro EVs (2K × g and 10K × g fractions) were initially isolated from hNSCs transduced with a lentiviral vector (LV) encoding PTBP1 isoform a–mCherry or mCherry alone. Whole Lysate and EV fractions were analyzed by Western blot for PTBP1 and flotillin-2. A specific PTBP1–mCherry band was detected in both the 2K × g and 10K × g fractions, whereas flotillin-2 was enriched in both EV fractions, particularly enriched in the 10K × g fractions. Notably, EVs contained multiple endogenous PTBP1 isoforms, including the canonical isoforms a and c (∼57 kDa), as well as additional low–molecular-weight isoforms (Figure 8A). These lower bands are consistent with protein-coding PTBP1 transcripts generated by alternative splicing, as revealed by RNA-seq analysis of control hNSCs (Figure 3A). Lysate and EVs derived from PTBP1–mCherry–transduced cells also contained an additional PTBP1 isoform (indicated by X), consistent with the data shown in Figure 3B. Importantly, PTBP1–mCherry levels in EVs were lower than those of endogenous PTBP1 isoforms, indicating that endogenous PTBP1 isoforms are preferentially and more efficiently loaded into EVs. Moreover, PTBP1–mCherry overexpression alters the overall PTBP1 isoform composition of EVs.

To visualize EV uptake, 10,000 × g EVs isolated from PTBP1–mCherry–expressing hNSCs were incubated with recipient PTBP1-KD hNSCs and analyzed by live holotomographic/fluorescence imaging under physiological hypoxic conditions. Live-cell imaging revealed efficient uptake of PTBP1–mCherry–containing EVs by recipient hNSCs (Figure 8B). Super-resolution confocal microscopy shows the presence of PTBP1–mCherry–positive signals within acceptor PTBP1-KD hNSCs, and quantitative confocal analysis revealed efficient delivery of PTBP1–mCherry to the nucleolus of recipient cells in EVs respect the control condition (Figure 8C).

Given the detrimental effects of PTBP1–mCherry overexpression on hNSC fate and on endogenous PTBP1 isoforms in both cell lysates and EVs (Figure 3 and Figure 8A), we next investigated whether EVs from untreated hNSCs could rescue the reduced proliferation observed in PTBP1KD cells (Figure 1E). EVs (2,000 × g and 10,000 × g fractions) were therefore isolated from untreated hNSCs, analyzed by Western blotting, and used for functional assays. High-resolution Western blot analysis showed that PTBP1 was present in both whole-cell lysates and EV fractions, whereas flotillin-2 was detected exclusively in EV fractions (Figure 8D). Importantly, EV fractions also contained lower–molecular-weight PTBP1 bands with apparent molecular weights consistent with distinct PTBP1 isoforms identified by RNA-seq analysis (Figure 3A). The PTBP1 banding pattern differed between the 2K × g and 10K × g fractions, indicating isoform-specific loading of PTBP1 into distinct EV populations.

We next investigated whether the nuclear transfer of PTBP1 observed in PTBP1 KD cells (Figure 8C) was sufficient to rescue the reduction in proliferation (Ki67+) induced by PTBP1 depletion (Figure 1E). EV fractions (2,000 xg and 10,000 xg) were then incubated with PTBP1KD cells while maintaining PTBP1 siRNA during the uptake experiment. After 72 h, cells were fixed and immunostained for Ki67, PTBP1, PTBP2, and Nestin. Quantitative analysis revealed that both EV fractions significantly restored Ki67 positivity in PTBP1KD acceptor cells compared with control conditions (Figure 8E). Super-resolution confocal microscopy further confirmed the presence of PTBP1-containing EVs associated with the surface of PTBP1KD cells (PTBP2-positive) (Figure 8F). Taken together, these data demonstrate that hNSCs produce EVs enriched in specific, previously uncharacterized PTBP1 isoforms, and that these EVs are efficiently captured by recipient hNSCs, thereby restoring proliferative capacity in PTBP1-depleted cells.

**Figure 8.**
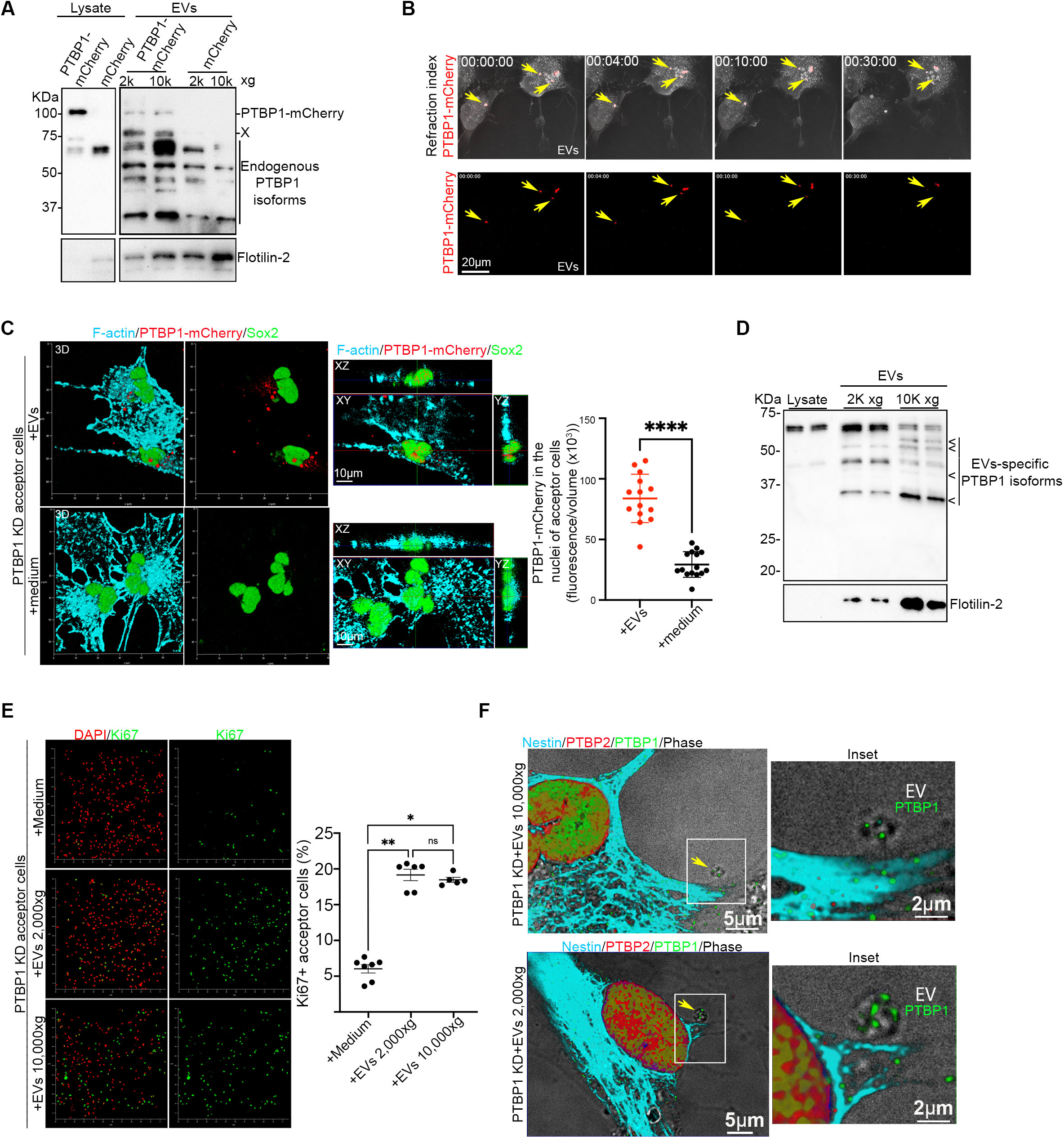
EVs Contain Specific PTBP1 Isoforms and EV Transfer Regulates Cell Proliferation in PTBP1-depleted cells. **A.** Representative Western blot analysis of cell lysates and EV fractions (2,000 × g and 10,000 × g) isolated from hNSCs transduced with a lentiviral vector (LV) encoding PTBP1 isoform a–mCherry or mCherry alone. EVs contain both PTBP1–mCherry and endogenous PTBP1. Notably, PTBP1–mCherry is inefficiently incorporated into EVs and alters the expression pattern of endogenous PTBP1 isoforms compared with mCherry alone (PTBP1 isoform indicated by X). Lanes corresponding to PTBP1 in cell lysates were exposed for a shorter time compared with all other lanes. (n = 3 for each group). **B.** Representative holotomographic/fluorescence time-lapse imaging of hNSCs incubated with EVs (10,000 × g fraction) derived from PTBP1–mCherry–expressing hNSCs shows efficient uptake of PTBP1–mCherry–containing EVs by recipient hNSC (n=3). **C.** Immunofluorescence analysis of PTBP1-KD cells incubated with the 10,000 × g EV fraction isolated from hNSCs transduced with a lentiviral vector encoding PTBP1 (NP_002810.1)–mCherry or with an equal volume of medium (control). Quantitative analysis of nuclear PTBP1–mCherry reveals efficient EV uptake and nuclear delivery of the recombinant protein. Each point represents the average nuclear PTBP1–mCherry fluorescence normalized to cell volume per random field (n = 3 for each group). **D.** Representative Western blot of total cell lysates and EV fractions (2,000 × g and 10,000 × g) isolated from normal hNSCs. Specific endogenous PTBP1 isoforms are detected in EVs, with distinct isoform distributions across fractions (n = 3 for each group). **E.** Analysis of Ki67 positivity in acceptor cells incubated with EV fractions (2,000 × g and 10,000 × g) isolated from normal hNSCs. Both fractions increase hNSC proliferation in PTBP1-depleted cells. Each point represents the percentage of Ki67-positive cells per random field (n = 3 for each group). **F.** Representative immunofluorescence analysis of PTBP1 (green), PTBP2 (red), and Nestin (cyan), followed by lightning super-resolution confocal microscopy, shows PTBP1-containing EVs (yellow arrows) associated with the surface of PTBP1-KD acceptor cells (PTBP2-positive). n = 3 independent experiments.

### PTBP1 is Highly Expressed in mouse V-SVZ and Localizes to the Nucleus, GFAP-positive processes, and F-actin TNT-like Structures Between NSCs

The cellular and subcellular localization of PTBP1 in the mouse brain has previously been characterized in the cortex ^66^, hippocampus and substantia nigra ^67–69^.

However, whether PTBP1 is expressed in the ventricular–subventricular zone (V-SVZ) and how it is distributed subcellularly within this neurogenic niche at level of NSCs has remained unknown.

To address this question, 20-µm-thick coronal sections from PD15 mouse brains were stained for PTBP1, GFAP, AQP4, and F-actin. Post-natal NSCs, B1-type and B2-type cells, were identified as GFAP-positive cells with cell bodies located either in the LV-SVZ and characterized by long, radially oriented GFAP-positive processes extending toward blood vessels. B1 cells establish direct contact with the lateral ventricle (LV), whereas B2 cells do not ^70–72^. AQP4 immunostaining was used to delineate the basal membrane of the ependymal layer and to identify astrocytic endfeet surrounding blood vessels.

Low-magnification (10×) reconstruction of entire coronal sections revealed PTBP1 expression in the cortex and striatum, with particularly strong enrichment in the V-SVZ (Figure 9A). Higher-magnification imaging of the lateral ventricle (Inset 1) showed that PTBP1 was especially enriched in the dorsal V-SVZ, wedge region, lateral wall, and ventral V-SVZ (Figure 9B).

Super-resolution imaging of the dorso-lateral V-SVZ (Inset 2) revealed high PTBP1 levels in the ependymal layer and in GFAP-positive cells extending long radial processes toward blood vessels, consistent with B1-type NSCs (Inset 3). A similar pattern was observed in GFAP-positive radial glia-like cells that did not contact the ependymal layer, consistent with B2-type NSCs. In both B1-type and B2-type NSCs, PTBP1 was detected in the nucleus and GFPA-positive processes (Insets 3 and 4, Figure 9C). Three-dimensional reconstruction and single-plane (XY1–XY4) analysis demonstrated that GFAP-positive processes extending from B1-type toward neighboring B2-type contained PTBP1 signals (Figure 9D, inset 5, red arrows). Furthermore, analysis of F-actin–positive processes and 3D reconstruction from serial confocal z-stacks acquired using super-resolution Lightning deconvolution microscopy revealed that B2-type cells are interconnected by thin F-actin positive intercellular protrusions resembling tunneling nanotubes (TNT-like structures) that contain PTBP1 (Figure 9E–F, inset 6, red arrows).

Together, these findings indicate that PTBP1 localizes not only to the nucleus and cytoplasm of NSCs but also to TNT-like structures connecting these cells. This observation is consistent with previous in vitro findings in human fetal NSCs and raises the possibility that PTBP1 may be transferred between NSCs via TNT-like structures in vivo.

**Figure 9.**
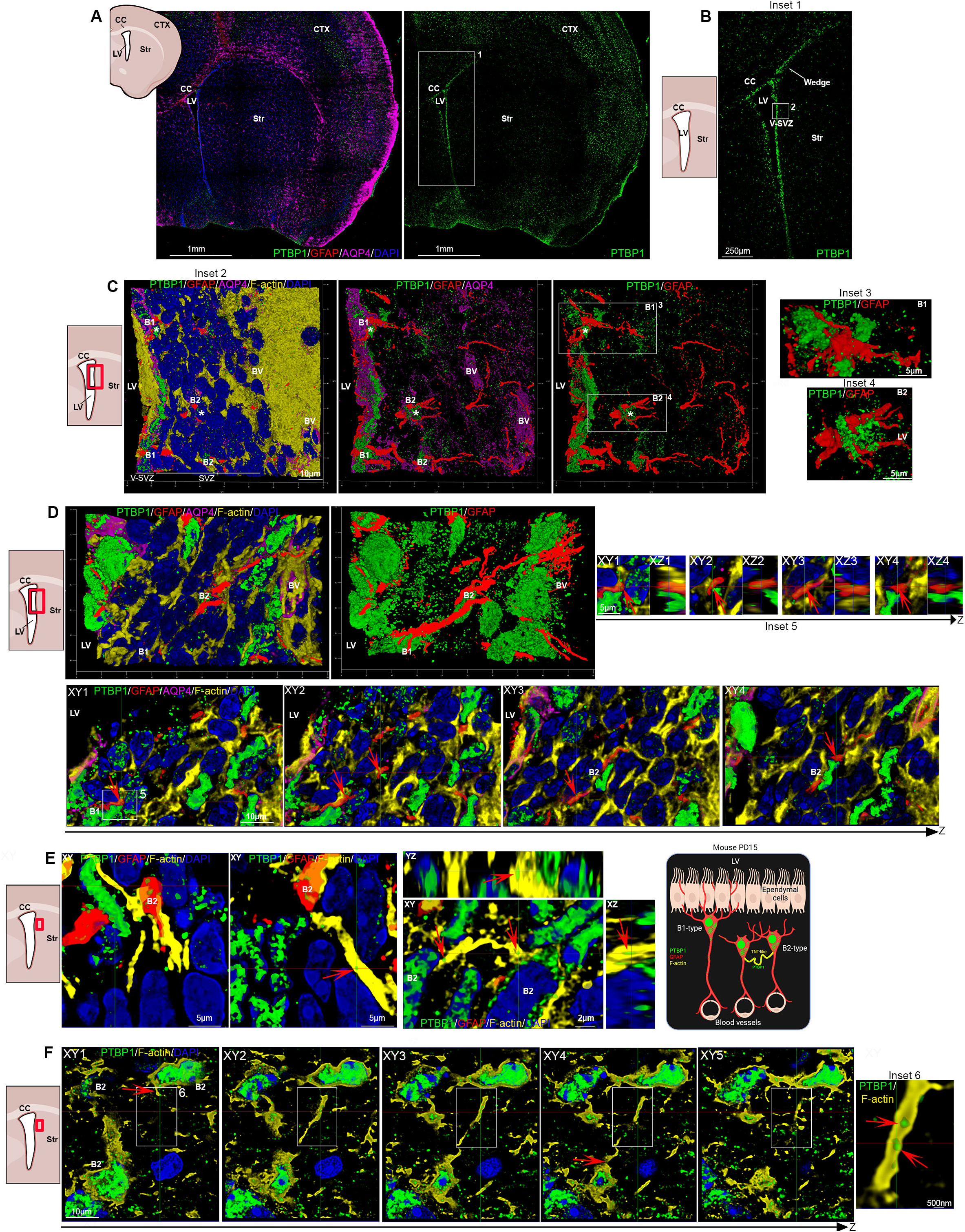
Subcellular localization of PTBP1 in the mouse V-SVZ reveals PTBP1 within TNT-like structures connecting NSCs. Immunofluorescence analysis of 20-µm coronal sections from PD15 mouse brains stained for PTBP1 (green), GFAP (red), F-actin (yellow), AQP4 (pink), and nuclei counterstained with DAPI (blue). (A) Low-magnification (10×) reconstruction of an entire coronal section showing prominent PTBP1 enrichment within the V-SVZ. (B) Higher-magnification view of the lateral ventricle illustrating strong PTBP1 signal along the dorsal V-SVZ, wedge region, lateral wall, and ventral V-SVZ. (C) Super-resolution imaging of the dorso-lateral V-SVZ showing PTBP1 expression in GFAP-NSCs, including B1-type contacting the ependymal layer and B2-type. PTBP1 is detected in both the nucleus and cytoplasm of B1- and B2-type NSCs (Insets 3–4). AQP4 staining delineates the ependymal basal membrane and astrocytic endfeet surrounding blood vessels. (D) Three-dimensional reconstruction and single-plane (XY1-4) analysis reveal PTBP1 signal within GFAP-positive radial processes extending between NSCs (Inset 5, red arrows). (E–F) Serial confocal z-stacks with Lightning super-resolution deconvolution demonstrate thin F-actin–positive intercellular bridges resembling tunneling nanotubes (TNT-like structures) connecting B2-type cells and containing PTBP1 (Inset 6, red arrows).

## Discussion

Here, we identify previously unrecognized roles for PTBP1 in the regulation of hNSCs. Our data demonstrate that PTBP1 is a key regulator of hNSC self-renewal and processes that critically influence neural stem cell fate, such as lipid droplet (LD) homeostasis, and mitochondrial dynamics^56,73,74^

Mitochondrial dynamics are now recognized as a central determinant of NSC fate, as highlighted in recent work (recently reviewed by by Garone et al. ^75^). Increasing evidence indicates that stem cell activation is associated with mitochondrial fission: daughter cells enriched in fragmented mitochondria tend to differentiate, whereas cells retaining fused mitochondria preferentially maintain stemness ^76–78^. In line with this paradigm, PTBP1 knockdown markedly impaired hNSC proliferation, increased mitochondrial number, and promoted mitochondrial fission, indicating that PTBP1 is required to preserve a mitochondria-associated stemness state. Although PTBP1 has recently been implicated in mitochondrial regulation ^76,79^ its role in mitochondrial dynamics in hNSCs had not previously been explored.

A similar knowledge gap existed for lipid metabolism. While lipid droplets and lipid metabolic pathways have emerged as important regulators of neural stem cell fate ^80–82^, our study is the first to demonstrate that PTBP1 directly influences LD homeostasis in hNSCs.

Importantly, our results indicate that hNSC fate is not governed solely by total PTBP1 abundance, but rather by the precise balance of PTBP1 isoforms. Alterations in isoform composition profoundly affected mitochondrial remodeling and lipid droplets, underscoring the need for tight isoform-specific regulation to maintain stemness. The reduction of endogenous PTBP1 isoforms (a and c) following ectopic overexpression of the isoform a likely reflects an autoregulatory mechanism that limits PTBP1 excess, analogous to what has been reported for the splicing factor hnRNPA1 ^83^.

Transcriptomic profiling further revealed that PTBP1 depletion induces widespread changes in the expression profile of hNSCs, including key regulators of neuronal differentiation such as PTBP2, FLNA, and GABBR1. While the PTBP1–PTBP2 axis and PTBP1-dependent repression of a poison exon in FLNA have been well characterized in mouse neural progenitors ^84^, our study provides the first evidence that these regulatory mechanisms operate in hNSCs. Similarly, PTBP1-mediated regulation of GABBR1, previously described only in a mouse catecholaminergic CNS cell line ^85^, is here extended to human NSCs.

Beyond protein-coding genes, we also uncovered a previously unappreciated role for PTBP1 in regulating the expression of long non-coding RNAs. The differential expression of some lncRNAs upon PTBP1 knockdown suggests additional layers of regulation whose contribution to hNSC fate remains to be investigated.

One of the most striking findings of this work is that PTBP1, a splicing factor that here we show to be essential for hNSCs stemness maintenance, also localizes to the cytoplasm and can be transferred between hNSCs via TNTs and EVs. These findings obtained from in vitro approaches were further supported by ex vivo analysis of mouse brain tissue, which for the first time provides evidence of TNT-like structures connecting B2-type NSCs in the post-natal juvenile mouse brain and demonstrates the presence of PTBP1 within both F-actin–positive TNT-like structures and GFAP-positive processes connecting NSCs.

To our knowledge, this represents the first demonstration of PTBP1 intercellular trafficking in hNSCs. Direct cell–cell contact promotes more rapid and efficient transfer when recipient cells were PTBP1-depleted, suggesting that TNT-mediated trafficking may function as a homeostatic mechanism to equilibrate PTBP1 levels within the stem cell pool. Notably, distinct PTBP1 isoforms were selectively enriched in EVs (large and micro EVs), supporting a functional role for EV-mediated transfer in shaping PTBP1 isoform composition and, consequently, the spatial regulation of hNSC fate. Although PTBP1 has been detected in EVs from other cell types ^86^, its presence in hNSC-derived EVs had not previously been reported.

Importantly, EV-mediated transfer of endogenous PTBP1 restored the proliferative capacity of PTBP1-depleted cells. This rescue did not occur through direct cell–cell contact, likely due to the concomitant transfer of PTBP2 from PTBP1-depleted cells to control cells, suggesting an additional layer of regulation in which PTBP2 itself acts as a cytosolic protein exchanged between hNSCs via TNTs.

These findings introduce a new conceptual framework for understanding how neural stem cells coordinate fate decisions within the subventricular zone niche. Because dynamic cell-to-cell communication is essential for NSC behavior and cerebral cortex development, PTBP1 trafficking via TNTs and EVs may represent a previously unrecognized mechanism contributing to brain development. Intriguingly, mutations affecting the nuclear–cytoplasmic transport of PTBP1 have recently been associated with neurodevelopmental disorders ^87^, highlighting the importance of PTBP1 cytosolic localization as a central mechanism in physiological neurogenesis, consistent with the intercellular transfer described here.

Importantly, we also show that hNSC-derived large and micro EVs contain specific, previously uncharacterized PTBP1 isoforms and can restore proliferation in PTBP1-depleted cells. This represents the first evidence linking PTBP1 isoforms in NSC-derived EVs to a functional rescue of stem cell proliferation. Although EVs from hNSCs have been widely explored in the context of neurodegenerative diseases, the molecular determinants of their biological activity remain poorly defined. Our findings suggest that PTBP1 isoforms may represent key functional components of EV-mediated effects, with potential implications for improving EV-based therapeutic strategies.

Finally, the demonstration that a splicing factor can traffic between cells via TNTs and EVs raises the intriguing possibility that intercellular PTBP1 transfer contributes to the conflicting results reported for astrocyte-to-neuron conversion following PTBP1 knockdown ^19,20^. While this approach has been proposed as a therapeutic strategy for neurodegenerative diseases, inconsistent outcomes across studies have fueled ongoing debate. We propose that PTBP1 intercellular transfer may represent an overlooked variable contributing to these discrepancies.

In conclusion, our study defines new functional roles for PTBP1 in human neural stem cells and reveals its dynamic intercellular transfer via TNTs and EVs, highlighting a novel mechanism contributing to the spatiotemporal regulation of neurogenesis. These findings provide new insight into the physiological control of neural stem cell behavior and may help elucidate how these processes become dysregulated in disease.

### Limitations of the study

Future studies will be required to determine whether hNSCs exchange PTBP1 in vivo, to elucidate the mechanisms responsible for the generation of alternative PTBP1 isoforms, and to define their role in hNSC fate. Characterizing TNTs and PTBP1 trafficking within rodent or primate SVZ niches, as well as localization of PTBP1 in fetal human brain tissue slices, will be essential. Advanced ultrastructural approaches—including EM-based methods applied to the human cerebellum ^88^ and 3D serial block-face SEM of the mouse V-SVZ ^89^ provide promising strategies to identify potential contact sites where TNT-mediated transfer of stemness-related signals may occur. However, in vivo tracking of PTBP1 remains technically challenging, as it would require multiphoton imaging, and PTBP1 fluorescent overexpression itself profoundly alters hNSC fate.

## Resource availability

### Lead contact

Further information and requests for data and reagents should be directed to and will be fulfilled by the lead contact Francesco Pisani.

## Materials availability

This study did not generate any unique reagents.

## Data availability

RNA-seq data have been deposited at Sequence Read Archive (SRA) (https://www.ncbi.nlm.nih.gov/sra) available as of the date of publication BioProject ID: PRJNA1425260. Microscopy and Western blotting data reported in this paper will be shared by the lead contact upon request. This paper does not report any original code or algorithms.

Any additional information required to reanalyze the data reported in this paper is available from the lead contact upon request.

## Ethics statement

The study was approved by the Ethical Committee of IRCCS Casa Sollievo della Sofferenza, Viale Cappuccini 1, San Giovanni Rotondo, 71,013 Foggia (PROT.01/CE25/01/12).

Mouse samples were kindly provided by Professor Giuseppe Calamita, University of Bari Aldo Moro (Authorization from the Italian Ministry of Health No. 158/2024-PR).

## Conflict of interes

None declared

## Acknowledgments

We thank ELIXIR-IT genomics infrastructure, through projects: PON CNR BIOMICS (PIR01_00017) and ELIXIRNextGenIT (Grant Code IR00000109). We thank Dr. Annunziata De Luisi for technical support. We thank Professor Giuseppe Calamita, Università degli studi di Bari aldo Moro, for providing the mice.

## Competing interests

The authors declare no competing interests.

Declaration of generative AI and AI-assisted technologies in the writing process:

During the preparation of this manuscript, the authors used ChatGPT as a language-editing tool to improve clarity, grammar, and readability.

## Author contributions

Conceptualization: F.P*^#^

Data curation: F.P.*, D.L.C, D.P.F., F.E, L.C., P.C., S.L. ^#^, D.A.M. ^#^

Formal analysis: F.P.*, D.L.C, D.P.F., D.P.F., F.E, L.C., P.C., S.L. ^#^, D.A.M. ^#^

Funding acquisition: F.P.*, P.E., P.G., V.A.L

Investigation: D.L.C, D.P.F., D.P.F., F.E, L.C., P.C., S.L., P.F., S.L. ^#^, D.A.M. ^#^

Methodology: P.D.C., G.M., S.L. ^#^, D.A.M. ^#^

Project administration: F.P.*, P.G., S.M.

Supervision: F.P.*, P.E., P.G., V.A.L., S.M., S.L. ^#^, D.A.M. ^#^

Validation: S.L. ^#^, D.A.M. ^#^

Writing – original draft: F.P.*, S.L. ^#^, D.A.M. ^#^

Writing – review & editing: F.P.*, D.L.C, D.P.F., F.E, L.C., P.C., S.L., P.F., P.D.C., G.M., P.E., P.G., V.A.L., S.M., S.L. ^#^, D.A.M. ^#^

*Lead contact

^#^ Equally contributing authors

## References

1. Conti L, Cattaneo E. Neural stem cell systems: physiological players or in vitro entities? Nat Rev Neurosci. 2010 Mar;11(3):176–87.

2. Ihunwo AO, Perego J, Martino G, Vicenzi E, Panina-Bordignon P. Neurogenesis and Viral Infection. Front Immunol. 2022;13:826091.

3. Liu JA, Cheung M. Neural crest stem cells and their potential therapeutic applications. Dev Biol. 2016 Nov 15;419(2):199–216.

4. Cipriani S, Ferrer I, Aronica E, Kovacs GG, Verney C, Nardelli J, et al. Hippocampal Radial Glial Subtypes and Their Neurogenic Potential in Human Fetuses and Healthy and Alzheimer’s Disease Adults. Cereb Cortex. 2018 Jul 1;28(7):2458–78.

5. Johansson PA, Dziegielewska KM, Liddelow SA, Saunders NR. The blood-CSF barrier explained: when development is not immaturity. Bioessays. 2008 Mar;30(3):237–48.

6. Lehtinen MK, Walsh CA. Neurogenesis at the brain-cerebrospinal fluid interface. Annu Rev Cell Dev Biol. 2011;27:653–79.

7. Yeh C, Li A, Chuang JZ, Saito M, Cáceres A, Sung CH. IGF-1 activates a cilium-localized noncanonical Gβγ signaling pathway that regulates cell-cycle progression. Dev Cell. 2013 Aug 26;26(4):358–68.

8. Jin Y, Liang X, Wang X. Alternative splicing in stem cells and development: research progress and emerging technologies. Cell Regen. 2025 Jun 4;14(1):20.

9. Nazim M, Lin CH, Feng AC, Xiao W, Yeom KH, Li M, et al. Alternative splicing of a chromatin modifier alters the transcriptional regulatory programs of stem cell maintenance and neuronal differentiation. Cell Stem Cell. 2024 May 2;31(5):754–771.e6.

10. Iannone C, Kainov Y, Zhuravskaya A, Hamid F, Nojima T, Makeyev E V. PTBP1-activated co-transcriptional splicing controls epigenetic status of pluripotent stem cells. Mol Cell. 2023 Jan 19;83(2):203–218.e9.

11. Vuong JK, Lin CH, Zhang M, Chen L, Black DL, Zheng S. PTBP1 and PTBP2 Serve Both Specific and Redundant Functions in Neuronal Pre-mRNA Splicing. Cell Rep. 2016 Dec 6;17(10):2766–75.

12. Keppetipola N, Sharma S, Li Q, Black DL. Neuronal regulation of pre-mRNA splicing by polypyrimidine tract binding proteins, PTBP1 and PTBP2. Crit Rev Biochem Mol Biol. 2012;47(4):360–78.

13. Boutz PL, Stoilov P, Li Q, Lin CH, Chawla G, Ostrow K, et al. A post-transcriptional regulatory switch in polypyrimidine tract-binding proteins reprograms alternative splicing in developing neurons. Genes Dev. 2007 Jul 1;21(13):1636–52.

14. Li Q, Zheng S, Han A, Lin CH, Stoilov P, Fu XD, et al. Correction: The splicing regulator PTBP2 controls a program of embryonic splicing required for neuronal maturation. Elife. 2017 Oct 9;6.

15. Ramos AD, Andersen RE, Liu SJ, Nowakowski TJ, Hong SJ, Gertz C, et al. The long noncoding RNA Pnky regulates neuronal differentiation of embryonic and postnatal neural stem cells. Cell Stem Cell. 2015 Apr 2;16(4):439–47.

16. Kamath R V, Leary DJ, Huang S. Nucleocytoplasmic shuttling of polypyrimidine tract-binding protein is uncoupled from RNA export. Mol Biol Cell. 2001 Dec;12(12):3808–20.

17. Cho CY, Chung SY, Lin S, Huang JS, Chen YL, Jiang SS, et al. PTBP1-mediated regulation of AXL mRNA stability plays a role in lung tumorigenesis. Sci Rep. 2019 Nov 15;9(1):16922.

18. Fan X, Zhao Z, Ma L, Huang X, Zhan Q, Song Y. PTBP1 promotes IRES-mediated translation of cyclin B1 in cancer. Acta Biochim Biophys Sin (Shanghai). 2022 May 25;54(5):696–707.

19. Hoang T, Kim DW, Appel H, Ozawa M, Zheng S, Kim J, et al. Ptbp1 deletion does not induce astrocyte-to-neuron conversion. Nature. 2023 Jun;618(7964):E1–7.

20. Hao Y, Hu J, Xue Y, Dowdy SF, Mobley WC, Qian H, et al. Reply to: Ptbp1 deletion does not induce astrocyte-to-neuron conversion. Nature. 2023 Jun;618(7964):E8–13.

21. Palese F, Rakotobe M, Zurzolo C. Transforming the concept of connectivity: unveiling tunneling nanotube biology and their roles in brain development and neurodegeneration. Physiol Rev. 2025 Jul 1;105(3):1823–65.

22. Rustom A, Saffrich R, Markovic I, Walther P, Gerdes HH. Nanotubular highways for intercellular organelle transport. Science. 2004 Feb 13;303(5660):1007–10.

23. Budinger D, Baker V, Heneka MT. Tunneling Nanotubes in the Brain. Results Probl Cell Differ. 2024;73:203–27.

24. Tarasiuk O, Scuteri A. Role of Tunneling Nanotubes in the Nervous System. Int J Mol Sci. 2022 Oct 19;23(20).

25. Neher JJ, Simons M. Protective lifelines: Tunneling nanotubes connect neurons and microglia. Neuron. 2024 Sep 25;112(18):2991–3.

26. Rostami J, Holmqvist S, Lindström V, Sigvardson J, Westermark GT, Ingelsson M, et al. Human Astrocytes Transfer Aggregated Alpha-Synuclein via Tunneling Nanotubes. J Neurosci. 2017 Dec 6;37(49):11835–53.

27. Errede M, Mangieri D, Longo G, Girolamo F, de Trizio I, Vimercati A, et al. Tunneling nanotubes evoke pericyte/endothelial communication during normal and tumoral angiogenesis. Fluids Barriers CNS. 2018 Oct 5;15(1):28.

28. Vargas JY, Loria F, Wu YJ, Córdova G, Nonaka T, Bellow S, et al. The Wnt/Ca2+ pathway is involved in interneuronal communication mediated by tunneling nanotubes. EMBO J. 2019 Dec 2;38(23):e101230.

29. Pisani F, Castagnola V, Simone L, Loiacono F, Svelto M, Benfenati F. Role of pericytes in blood-brain barrier preservation during ischemia through tunneling nanotubes. Cell Death Dis. 2022 Jul 5;13(7):582.

30. Scheiblich H, Eikens F, Wischhof L, Opitz S, Jüngling K, Cserép C, et al. Microglia rescue neurons from aggregate-induced neuronal dysfunction and death through tunneling nanotubes. Neuron. 2024 Sep 25;112(18):3106–3125.e8.

31. Capobianco DL, Simone L, Svelto M, Pisani F. Intercellular crosstalk mediated by tunneling nanotubes between central nervous system cells. What we need to advance. Front Physiol. 2023;14:1214210.

32. Capobianco DL, De Zio R, Profico DC, Gelati M, Simone L, D’Erchia AM, et al. Human neural stem cells derived from fetal human brain communicate with each other and rescue ischemic neuronal cells through tunneling nanotubes. Cell Death Dis. 2024 Sep 1;15(8):639.

33. Simone L, Capobianco DL, Di Palma F, Binda E, Legnani FG, Vescovi AL, et al. GFAP serves as a structural element of tunneling nanotubes between glioblastoma cells and could play a role in the intercellular transfer of mitochondria. Front Cell Dev Biol. 2023;11:1221671.

34. Chang M, Krüssel S, Parajuli LK, Kim J, Lee D, Merodio A, et al. Intercellular communication in the brain through a dendritic nanotubular network. Science. 2025 Oct 2;390(6768):eadr7403.

35. van Niel G, D’Angelo G, Raposo G. Shedding light on the cell biology of extracellular vesicles. Nat Rev Mol Cell Biol. 2018 Apr;19(4):213–28.

36. Jiang Y, Liu X, Ye J, Ma Y, Mao J, Feng D, et al. Migrasomes, a new mode of intercellular communication. Cell Commun Signal. 2023 May 8;21(1):105.

37. Santavanond JP, Rutter SF, Atkin-Smith GK, Poon IKH. Apoptotic Bodies: Mechanism of Formation, Isolation and Functional Relevance. Subcell Biochem. 2021;97:61–88.

38. Zhu Z, Zhang Q, Feng J, Zebaze Dongmo S, Zhang Q, Huang S, et al. Neural Stem Cell-Derived Small Extracellular Vesicles: key Players in Ischemic Stroke Therapy - A Comprehensive Literature Review. Int J Nanomedicine. 2024;19:4279–95.

39. Spinelli M, Fusco S, Grassi C. Therapeutic potential of stem cell-derived extracellular vesicles in neurodegenerative diseases associated with cognitive decline. Stem Cells. 2025 Feb 12;43(2).

40. Wang M, Chen D, Pan R, Sun Y, He X, Qiu Y, et al. Neural stem cell-derived small extracellular vesicles: a new therapy approach in neurological diseases. Front Immunol. 2025;16:1548206.

41. Derkus B, Isik M, Eylem CC, Ergin I, Camci CB, Bilgin S, et al. Xenogenic Neural Stem Cell-Derived Extracellular Nanovesicles Modulate Human Mesenchymal Stem Cell Fate and Reconstruct Metabolomic Structure. Adv Biol. 2022 Jun;6(6):e2101317.

42. Sun X, Chen Y, Zhang Y, Cheng T, Peng H, Sun Y, et al. Exosomes released from immature neurons regulate adult neural stem cell differentiation through microRNA-7a-5p. Stem Cells. 2025 Feb 12;43(2).

43. Ewels P, Magnusson M, Lundin S, Käller M. MultiQC: summarize analysis results for multiple tools and samples in a single report. Bioinformatics. 2016 Oct 1;32(19):3047–8.

44. Chen S, Zhou Y, Chen Y, Gu J. fastp: an ultra-fast all-in-one FASTQ preprocessor. Bioinformatics. 2018 Sep 1;34(17):i884–90.

45. Dobin A, Davis CA, Schlesinger F, Drenkow J, Zaleski C, Jha S, et al. STAR: ultrafast universal RNA-seq aligner. Bioinformatics. 2013 Jan 1;29(1):15–21.

46. Liao Y, Smyth GK, Shi W. featureCounts: an efficient general purpose program for assigning sequence reads to genomic features. Bioinformatics. 2014 Apr 1;30(7):923–30.

47. Love MI, Huber W, Anders S. Moderated estimation of fold change and dispersion for RNA-seq data with DESeq2. Genome Biol. 2014;15(12):550.

48. Bray NL, Pimentel H, Melsted P, Pachter L. Near-optimal probabilistic RNA-seq quantification. Nat Biotechnol. 2016 May 4;34(5):525–7.

49. Shen S, Park JW, Lu Z xiang, Lin L, Henry MD, Wu YN, et al. rMATS: robust and flexible detection of differential alternative splicing from replicate RNA-Seq data. Proc Natl Acad Sci U S A. 2014 Dec 23;111(51):E5593–601.

50. Filomena E, Picardi E, Pesole G, D’Erchia AM. IsoPrimer: a pipeline for designing isoform-aware primer pairs for comprehensive gene expression quantification. Bioinformatics advances. 2025;5(1):vbaf171.

51. Leone MA, Gelati M, Profico DC, Gobbi C, Pravatà E, Copetti M, et al. Phase I clinical trial of intracerebroventricular transplantation of allogeneic neural stem cells in people with progressive multiple sclerosis. Cell Stem Cell. 2023 Dec 7;30(12):1597–1609.e8.

52. Vuong CK, Black DL, Zheng S. The neurogenetics of alternative splicing. Nat Rev Neurosci. 2016 May;17(5):265–81.

53. Wang Y, Barthez M, Chen D. Mitochondrial regulation in stem cells. Trends Cell Biol. 2024 Aug;34(8):685–94.

54. Ramosaj M, Madsen S, Maillard V, Scandella V, Sudria-Lopez D, Yuizumi N, et al. Lipid droplet availability affects neural stem/progenitor cell metabolism and proliferation. Nat Commun. 2021 Dec 21;12(1):7362.

55. Boutz PL, Stoilov P, Li Q, Lin CH, Chawla G, Ostrow K, et al. A post-transcriptional regulatory switch in polypyrimidine tract-binding proteins reprograms alternative splicing in developing neurons. Genes Dev. 2007 Jul 1;21(13):1636–52.

56. Makeyev E V, Zhang J, Carrasco MA, Maniatis T. The MicroRNA miR-124 promotes neuronal differentiation by triggering brain-specific alternative pre-mRNA splicing. Mol Cell. 2007 Aug 3;27(3):435–48.

57. Zhang X, Chen MH, Wu X, Kodani A, Fan J, Doan R, et al. Cell-Type-Specific Alternative Splicing Governs Cell Fate in the Developing Cerebral Cortex. Cell. 2016 Aug 25;166(5):1147–1162.e15.

58. Vuong CK, Black DL, Zheng S. The neurogenetics of alternative splicing. Nat Rev Neurosci. 2016 May;17(5):265–81.

59. Su CH, D D, Tarn WY. Alternative Splicing in Neurogenesis and Brain Development. Front Mol Biosci. 2018;5:12.

60. Pin JP, Bettler B. Organization and functions of mGlu and GABAB receptor complexes. Nature. 2016 Dec 1;540(7631):60–8.

61. Lian G, Wong T, Lu J, Hu J, Zhang J, Sheen V. Cytoskeletal Associated Filamin A and RhoA Affect Neural Progenitor Specification During Mitosis. Cereb Cortex. 2019 Mar 1;29(3):1280–90.

62. Lian G, Lu J, Hu J, Zhang J, Cross SH, Ferland RJ, et al. Filamin a regulates neural progenitor proliferation and cortical size through Wee1-dependent Cdk1 phosphorylation. J Neurosci. 2012 May 30;32(22):7672–84.

63. Falace A, Corbieres L, Palminha C, Guarnieri FC, Schaller F, Buhler E, et al. FLNA regulates neuronal maturation by modulating RAC1-Cofilin activity in the developing cortex. Neurobiol Dis. 2024 Aug;198:106558.

64. Zhang X, Chen MH, Wu X, Kodani A, Fan J, Doan R, et al. Cell-Type-Specific Alternative Splicing Governs Cell Fate in the Developing Cerebral Cortex. Cell. 2016 Aug;166(5):1147–1162.e15.

65. Wang Y, Barthez M, Chen D. Mitochondrial regulation in stem cells. Trends Cell Biol. 2024 Aug;34(8):685–94.

66. Scandella V, Petrelli F, Moore DL, Braun SMG, Knobloch M. Neural stem cell metabolism revisited: a critical role for mitochondria. Trends Endocrinol Metab. 2023 Aug;34(8):446–61.

67. Garone C, De Giorgio F, Carli S. Mitochondrial metabolism in neural stem cells and implications for neurodevelopmental and neurodegenerative diseases. J Transl Med. 2024 Mar 4;22(1):238.

68. Khacho M, Clark A, Svoboda DS, Azzi J, MacLaurin JG, Meghaizel C, et al. Mitochondrial Dynamics Impacts Stem Cell Identity and Fate Decisions by Regulating a Nuclear Transcriptional Program. Cell Stem Cell. 2016 Aug 4;19(2):232–47.

69. Iwata R, Casimir P, Vanderhaeghen P. Mitochondrial dynamics in postmitotic cells regulate neurogenesis. Science. 2020 Aug 14;369(6505):858–62.

70. Baker N, Wade S, Triolo M, Girgis J, Chwastek D, Larrigan S, et al. The mitochondrial protein OPA1 regulates the quiescent state of adult muscle stem cells. Cell Stem Cell. 2022 Sep 1;29(9):1315–1332.e9.

71. Yang Y, Wen H, Li Y, Zeng X, Wei G, Gu Z, et al. Cellular senescence induced by down-regulation of PTBP1 correlates with exon skipping of mitochondrial-related gene NDUFV3. Life medicine. 2024 Apr;3(2):lnae021.

72. Wang C, Zhang H, Fan J, Li Q, Guo R, Pan J, et al. Inhibition of integrated stress response protects against lipid-induced senescence in hypothalamic neural stem cells in adamantinomatous craniopharyngioma. Neuro Oncol. 2023 Apr 6;25(4):720–32.

73. Knobloch M. The Role of Lipid Metabolism for Neural Stem Cell Regulation. Brain Plast. 2017 Nov 9;3(1):61–71.

74. Ramosaj M, Madsen S, Maillard V, Scandella V, Sudria-Lopez D, Yuizumi N, et al. Lipid droplet availability affects neural stem/progenitor cell metabolism and proliferation. Nat Commun. 2021 Dec 21;12(1):7362.

75. Suzuki H, Matsuoka M. hnRNPA1 autoregulates its own mRNA expression to remain non-cytotoxic. Mol Cell Biochem. 2017 Mar;427(1–2):123–31.

76. Zhang X, Chen MH, Wu X, Kodani A, Fan J, Doan R, et al. Cell-Type-Specific Alternative Splicing Governs Cell Fate in the Developing Cerebral Cortex. Cell. 2016 Aug 25;166(5):1147–1162.e15.

77. Makeyev E V, Zhang J, Carrasco MA, Maniatis T. The MicroRNA miR-124 promotes neuronal differentiation by triggering brain-specific alternative pre-mRNA splicing. Mol Cell. 2007 Aug 3;27(3):435–48.

78. Gao Y, Zhang Y, Mi N, Miao W, Zhang J, Liu Y, et al. Exploring the link between M1 macrophages and EMT of amniotic epithelial cells: implications for premature rupture of membranes. J Nanobiotechnology. 2025 Mar 4;23(1):163.

79. Masson A, Paccaud J, Orefice M, Colin E, Mäkitie O, Cormier-Daire V, et al. PTBP1 variants displaying altered nucleocytoplasmic distribution are responsible for a neurodevelopmental disorder with skeletal dysplasia. J Clin Invest. 2025 Nov 17;135(22).

80. Cordero Cervantes D, Khare H, Wilson AM, Mendoza ND, Coulon-Mahdi O, Lichtman JW, et al. 3D reconstruction of the cerebellar germinal layer reveals tunneling connections between developing granule cells. Sci Adv. 2023 Apr 5;9(14):eadf3471.

81. Takemura S, Kawase K, Wolbeck L, Nakamura Y, Matsumoto M, Jinnou H, et al. Transformation of radial glia into postnatal neural stem cells depends on birth. Cell Rep. 2025 Aug 26;44(8):116029.

