## Supplementary Table 1 for "Intercellular Transfer of PTBP1 Drives Human Neural Stem Cell Fate"

| ens_gene | ext_gene | gene_type | log2FoldChange | pvalue | padj | average_treated | average_control | siRNAPTBP1_1 | siRNAPTBP1_2 | siRNAPTBP1_3 | siRNActrl2 | siRNActrl1 | siRNActrl3 |
|---|---|---|---|---|---|---|---|---|---|---|---|---|---|
| ENSG00000204681 | GABBR1 | protein_coding | 2,233 | 1.500e-69 | 1.418e-65 | 9840,98 | 2094,55 | 9427,568 | 9360,141 | 10735,243 | 1978,163 | 2224,11 | 2081,392 |
| ENSG00000117569 | PTBP2 | protein_coding | 2,065 | 1.806e-51 | 8.536e-48 | 12979,3 | 3099,25 | 12889,171 | 12166,665 | 13882,052 | 3434,097 | 2663,48 | 3200,176 |
| ENSG00000259916 | AQP7B | protein_coding | 1,82 | 2.821e-05 | 5.178e-03 | 57,08 | 16,76 | 59,65 | 62,526 | 49,059 | 23,002 | 17,639 | 9,641 |
| ENSG00000176125 | UFSP1 | protein_coding | 1,579 | 1.925e-05 | 3.926e-03 | 77,41 | 25,89 | 67,225 | 107,187 | 57,82 | 24,706 | 24,053 | 28,922 |
| ENSG00000007516 | BAIAP3 | protein_coding | 1,484 | 1.668e-13 | 2.253e-10 | 3935,94 | 1410,4 | 3644,341 | 5078,004 | 3085,484 | 1491,715 | 1704,563 | 1034,912 |
| ENSG00000285935 | ENSG00000285935 | lncRNA | 1,443 | 1.648e-05 | 3.500e-03 | 79,61 | 29,27 | 92,789 | 77,711 | 68,333 | 30,669 | 27,26 | 29,886 |
| ENSG00000169840 | GSX1 | protein_coding | 1,345 | 1.158e-10 | 1.152e-07 | 1611,12 | 635,64 | 1387,103 | 2270,587 | 1175,673 | 565,676 | 675,09 | 666,161 |
| ENSG00000265401 | ENSG00000265401 | lncRNA | 1,297 | 5.531e-05 | 8.367e-03 | 106,91 | 42,94 | 139,184 | 100,935 | 80,598 | 51,115 | 30,467 | 47,239 |
| ENSG00000272114 | ENSG00000272114 | lncRNA | 1,266 | 1.747e-04 | 1.878e-02 | 156,62 | 65,64 | 234,813 | 105,401 | 129,657 | 91,156 | 60,935 | 44,829 |
| ENSG00000297142 | ENSG00000297142 | lncRNA | 1,239 | 2.792e-05 | 5.176e-03 | 117,16 | 49,11 | 118,354 | 142,023 | 91,11 | 49,411 | 40,088 | 57,843 |
| ENSG00000304919 | ENSG00000304919 | lncRNA | 1,213 | 5.189e-05 | 7.978e-03 | 208,05 | 92,19 | 154,333 | 285,833 | 183,973 | 75,821 | 134,697 | 66,038 |
| ENSG00000158555 | GDPD5 | protein_coding | 1,157 | 5.486e-10 | 4.716e-07 | 2734,82 | 1225,76 | 2282,803 | 3391,588 | 2530,062 | 976,303 | 1189,826 | 1511,154 |
| ENSG00000106701 | FSD1L | protein_coding | 1,147 | 8.357e-18 | 1.436e-14 | 4001,87 | 1803,38 | 3838,441 | 3856,96 | 4310,217 | 1932,159 | 1606,747 | 1871,228 |
| ENSG00000171798 | KNDC1 | protein_coding | 1,131 | 7.852e-09 | 5.120e-06 | 310,39 | 142,68 | 291,623 | 320,669 | 318,886 | 124,381 | 161,958 | 141,716 |
| ENSG00000299828 | ENSG00000299828 | lncRNA | 1,125 | 3.562e-05 | 6.124e-03 | 155,35 | 70,69 | 162,854 | 185,792 | 117,392 | 63,894 | 60,935 | 87,247 |
| ENSG00000149591 | TAGLN | protein_coding | 1,089 | 9.699e-10 | 7.974e-07 | 2463,94 | 1155,93 | 2338,665 | 2281,305 | 2771,855 | 879,183 | 1135,306 | 1453,311 |
| ENSG00000188818 | ZDHHC11 | protein_coding | 1,058 | 6.709e-05 | 9.539e-03 | 132,5 | 64,7 | 147,705 | 142,917 | 106,879 | 67,302 | 68,952 | 57,843 |
| ENSG00000124942 | AHNAK | protein_coding | 1,037 | 3.391e-10 | 3.053e-07 | 7287,56 | 3548,42 | 7123,935 | 8199,837 | 6538,914 | 3103,552 | 3082,003 | 4459,713 |
| ENSG00000300552 | ENSG00000300552 | lncRNA | 1,029 | 1.055e-04 | 1.355e-02 | 290,16 | 145,36 | 267,006 | 312,63 | 290,852 | 137,159 | 211,667 | 87,247 |
| ENSG00000259952 | ENSG00000259952 | lncRNA | 1,022 | 2.626e-04 | 2.548e-02 | 150,2 | 75,3 | 167,589 | 184,898 | 98,119 | 81,785 | 78,573 | 65,556 |
| ENSG00000138769 | CDKL2 | protein_coding | 1,02 | 9.625e-06 | 2.247e-03 | 209,3 | 103,31 | 185,578 | 223,307 | 219,015 | 124,381 | 97,816 | 87,729 |
| ENSG00000110013 | SIAE | protein_coding | 1,014 | 1.407e-11 | 1.774e-08 | 6056,22 | 2994,35 | 5684,756 | 5654,136 | 6829,766 | 2721,039 | 2719,603 | 3542,416 |
| ENSG00000295972 | ENSG00000295972 | lncRNA | 0,964 | 1.060e-04 | 1.355e-02 | 171,96 | 89,11 | 168,535 | 203,656 | 143,674 | 103,934 | 88,195 | 75,196 |
| ENSG00000054803 | CBLN4 | protein_coding | 0,952 | 1.016e-06 | 3.431e-04 | 492,13 | 253,92 | 398,615 | 604,716 | 473,073 | 234,279 | 246,945 | 280,54 |
| ENSG00000183979 | NPB | protein_coding | 0,944 | 4.533e-04 | 3.779e-02 | 131,12 | 69,57 | 123,088 | 159,888 | 110,384 | 67,302 | 80,177 | 61,217 |
| ENSG00000148204 | CRB2 | protein_coding | 0,943 | 1.279e-06 | 4.244e-04 | 2725,7 | 1420,29 | 2194,748 | 3632,76 | 2349,594 | 1203,766 | 1714,184 | 1342,927 |
| ENSG00000276564 | ENSG00000276564 | lncRNA | 0,937 | 6.290e-07 | 2.379e-04 | 502,82 | 259,86 | 460,158 | 438,575 | 609,738 | 270,059 | 226,099 | 283,432 |
| ENSG00000130822 | PNCK | protein_coding | 0,923 | 2.396e-04 | 2.372e-02 | 211,3 | 114,04 | 207,355 | 223,307 | 203,246 | 115,861 | 149,129 | 77,124 |
| ENSG00000187122 | SLIT1 | protein_coding | 0,916 | 6.500e-04 | 4.745e-02 | 213,84 | 116,29 | 203,568 | 224,2 | 213,759 | 99,675 | 171,579 | 77,606 |
| ENSG00000139187 | KLRG1 | protein_coding | 0,901 | 2.566e-05 | 4.852e-03 | 205,48 | 110,82 | 208,302 | 222,414 | 185,725 | 111,602 | 113,851 | 107,01 |
| ENSG00000185567 | AHNAK2 | protein_coding | 0,894 | 3.826e-04 | 3.412e-02 | 225,74 | 120,72 | 190,312 | 173,286 | 313,63 | 109,046 | 133,094 | 120,025 |
| ENSG00000101335 | MYL9 | protein_coding | 0,888 | 9.775e-07 | 3.381e-04 | 546,31 | 295,79 | 592,714 | 466,265 | 579,952 | 247,909 | 331,933 | 307,533 |
| ENSG00000183696 | UPP1 | protein_coding | 0,882 | 1.931e-05 | 3.926e-03 | 289,34 | 156,01 | 240,494 | 333,174 | 294,356 | 151,642 | 144,319 | 172,084 |
| ENSG00000148411 | NACC2 | protein_coding | 0,877 | 2.101e-07 | 9.239e-05 | 22805,77 | 12422,34 | 19251,856 | 29448,848 | 19716,614 | 11053,687 | 13025,553 | 13187,774 |
| ENSG00000140859 | KIFC3 | protein_coding | 0,872 | 1.973e-06 | 6.047e-04 | 1272,26 | 698,56 | 1162,705 | 1620,316 | 1033,751 | 637,238 | 808,184 | 650,254 |
| ENSG00000229124 | VIM-AS1 | lncRNA | 0,868 | 1.954e-08 | 1.192e-05 | 3291,54 | 1803,67 | 3800,568 | 3298,693 | 2775,359 | 2047,168 | 1627,593 | 1736,261 |
| ENSG00000130054 | NALF2 | protein_coding | 0,83 | 6.275e-04 | 4.599e-02 | 248,8 | 141,34 | 255,644 | 282,26 | 208,502 | 182,311 | 139,508 | 102,19 |
| ENSG00000197461 | PDGFA | protein_coding | 0,824 | 4.335e-08 | 2.242e-05 | 1946,76 | 1099,49 | 1691,035 | 2108,019 | 2041,221 | 954,153 | 1175,395 | 1168,915 |
| ENSG00000143847 | PPFIA4 | protein_coding | 0,819 | 2.025e-06 | 6.047e-04 | 5664,37 | 3214,84 | 6518,912 | 4559,038 | 5915,159 | 3089,069 | 3901,412 | 2654,04 |
| ENSG00000123095 | BHLHE41 | protein_coding | 0,813 | 3.500e-08 | 1.996e-05 | 3395,98 | 1935,29 | 3525,041 | 3032,511 | 3630,394 | 1950,049 | 2200,057 | 1655,762 |
| ENSG00000113212 | PCDHB7 | protein_coding | 0,812 | 6.804e-04 | 4.892e-02 | 185,63 | 107,24 | 173,27 | 222,414 | 161,195 | 117,565 | 115,455 | 88,693 |
| ENSG00000049769 | PPP1R3F | protein_coding | 0,81 | 4.038e-05 | 6.756e-03 | 1743,65 | 998,97 | 1623,81 | 2072,29 | 1534,858 | 965,228 | 1305,281 | 726,415 |
| ENSG00000167995 | BEST1 | protein_coding | 0,81 | 3.100e-04 | 2.873e-02 | 239,27 | 136,52 | 299,198 | 220,627 | 197,99 | 150,79 | 121,869 | 136,896 |
| ENSG00000267322 | SNHG22 | lncRNA | 0,804 | 5.595e-05 | 8.396e-03 | 729,49 | 420,08 | 789,655 | 897,695 | 501,107 | 436,184 | 429,749 | 394,298 |
| ENSG00000129654 | FOXJ1 | protein_coding | 0,803 | 4.134e-05 | 6.856e-03 | 619,21 | 353,83 | 586,087 | 749,419 | 522,132 | 310,951 | 314,294 | 436,234 |
| ENSG00000184156 | KCNQ3 | protein_coding | 0,793 | 2.169e-06 | 6.311e-04 | 1258,88 | 727,76 | 1180,695 | 1103,137 | 1492,807 | 709,651 | 840,255 | 633,383 |
| ENSG00000197444 | OGDHL | protein_coding | 0,785 | 2.140e-05 | 4.172e-03 | 1459,74 | 850,65 | 1377,635 | 1377,358 | 1624,216 | 821,253 | 1092,011 | 638,686 |
| ENSG00000198467 | TPM2 | protein_coding | 0,77 | 6.182e-06 | 1.518e-03 | 5077,03 | 2980,76 | 5675,288 | 5546,949 | 4008,852 | 3181,076 | 3357,812 | 2403,386 |
| ENSG00000127540 | UQCR11 | protein_coding | 0,768 | 5.247e-04 | 4.033e-02 | 5851,83 | 3439,33 | 5845,717 | 8040,842 | 3668,941 | 4176,121 | 3336,966 | 2804,914 |
| ENSG00000163013 | FBXO41 | protein_coding | 0,757 | 3.238e-05 | 5.776e-03 | 1983,05 | 1177,59 | 1678,726 | 2258,975 | 2011,434 | 1068,31 | 1510,534 | 953,931 |
| ENSG00000219438 | TAFA5 | protein_coding | 0,741 | 2.487e-05 | 4.751e-03 | 2860,34 | 1709,82 | 2561,17 | 3577,38 | 2442,456 | 1557,313 | 1524,966 | 2047,168 |
| ENSG00000124313 | IQSEC2 | protein_coding | 0,739 | 2.675e-05 | 5.008e-03 | 385,7 | 231,28 | 356,007 | 429,643 | 371,45 | 234,279 | 227,703 | 231,855 |
| ENSG00000080573 | COL5A3 | protein_coding | 0,739 | 8.165e-05 | 1.126e-02 | 2911,43 | 1746,85 | 2988,19 | 3738,161 | 2007,93 | 1719,178 | 1832,846 | 1688,54 |
| ENSG00000250132 | ENSG00000250132 | lncRNA | 0,737 | 4.812e-05 | 7.459e-03 | 367,05 | 222,18 | 400,508 | 353,718 | 346,92 | 215,536 | 248,549 | 202,451 |
| ENSG00000204347 | BTBD17 | protein_coding | 0,731 | 2.487e-04 | 2.437e-02 | 6050,26 | 3649,6 | 4725,619 | 8317,743 | 5107,432 | 3216,005 | 4419,356 | 3313,453 |
| ENSG00000187024 | PTRH1 | protein_coding | 0,721 | 7.727e-05 | 1.090e-02 | 503,32 | 307,03 | 446,903 | 612,755 | 450,295 | 309,248 | 325,519 | 286,324 |
| ENSG00000196557 | CACNA1H | protein_coding | 0,719 | 1.078e-05 | 2.456e-03 | 6945,45 | 4224,06 | 6436,537 | 6883,218 | 7516,598 | 3859,206 | 5331,77 | 3481,198 |
| ENSG00000305069 | ENSG00000305069 | lncRNA | 0,718 | 5.950e-04 | 4.447e-02 | 242,82 | 146,28 | 251,856 | 229,56 | 247,049 | 129,492 | 136,301 | 173,048 |
| ENSG00000196581 | AJAP1 | protein_coding | 0,713 | 4.402e-05 | 7.172e-03 | 4774,59 | 2909,85 | 4607,266 | 5567,493 | 4149,022 | 2563,433 | 2464,641 | 3701,484 |
| ENSG00000100065 | CARD10 | protein_coding | 0,709 | 6.591e-04 | 4.787e-02 | 372,35 | 228,78 | 311,506 | 434,109 | 371,45 | 177,2 | 266,188 | 242,942 |
| ENSG00000087085 | ACHE | protein_coding | 0,706 | 1.199e-04 | 1.462e-02 | 1086,46 | 669,51 | 997,957 | 1064,728 | 1196,698 | 618,495 | 862,704 | 527,337 |
| ENSG00000103154 | NECAB2 | protein_coding | 0,695 | 8.125e-05 | 1.126e-02 | 439,65 | 273,88 | 415,658 | 460,013 | 443,287 | 266,652 | 315,897 | 239,085 |
| ENSG00000170439 | TMT1B | protein_coding | 0,694 | 3.391e-05 | 5.883e-03 | 2308,68 | 1423,66 | 2476,902 | 2258,975 | 2190,151 | 1248,066 | 1213,88 | 1809,047 |
| ENSG00000132970 | WASF3 | protein_coding | 0,692 | 1.555e-08 | 9.799e-06 | 29542,57 | 18281,45 | 29506,951 | 27703,48 | 31417,275 | 17666,305 | 17488,204 | 19689,834 |
| ENSG00000099365 | STX1B | protein_coding | 0,692 | 3.820e-08 | 2.064e-05 | 4474,39 | 2772,44 | 4499,327 | 4436,666 | 4487,181 | 2786,637 | 3003,43 | 2527,267 |
| ENSG00000157542 | KCNJ6 | protein_coding | 0,691 | 2.399e-06 | 6.873e-04 | 3377,71 | 2087,02 | 3310,111 | 3204,904 | 3618,129 | 2056,539 | 1775,118 | 2429,416 |
| ENSG00000120913 | PDLIM2 | protein_coding | 0,686 | 1.950e-04 | 2.060e-02 | 1422,96 | 887,59 | 1160,811 | 1767,699 | 1340,372 | 763,322 | 1072,768 | 826,676 |
| ENSG00000273032 | DGCR5 | lncRNA | 0,685 | 6.214e-05 | 9.109e-03 | 708,91 | 441,3 | 710,121 | 775,322 | 641,276 | 364,623 | 457,009 | 502,272 |
| ENSG00000105404 | RABAC1 | protein_coding | 0,685 | 2.810e-04 | 2.657e-02 | 3268,39 | 2036,4 | 2740,121 | 4265,166 | 2799,889 | 1894,674 | 2480,676 | 1733,851 |
| ENSG00000127528 | KLF2 | protein_coding | 0,684 | 3.272e-05 | 5.781e-03 | 506,05 | 316,15 | 494,244 | 536,83 | 487,09 | 340,769 | 322,311 | 285,36 |
| ENSG00000006015 | REX1BD | protein_coding | 0,678 | 4.723e-04 | 3.862e-02 | 1462,99 | 917,15 | 1345,443 | 1964,209 | 1079,306 | 929,447 | 986,177 | 835,835 |
| ENSG00000107317 | PTGDS | protein_coding | 0,676 | 5.822e-05 | 8.668e-03 | 9384,74 | 5878,3 | 9931,28 | 9663,838 | 8559,109 | 7007,058 | 6274,651 | 4353,185 |
| ENSG00000128564 | VGF | protein_coding | 0,676 | 1.748e-04 | 1.878e-02 | 872 | 547,83 | 740,42 | 862,859 | 1012,726 | 510,301 | 667,073 | 466,12 |
| ENSG00000260630 | SNAI3-AS1 | lncRNA | 0,675 | 1.119e-04 | 1.402e-02 | 570,42 | 359,91 | 518,862 | 642,231 | 550,166 | 361,215 | 410,506 | 308,015 |
| ENSG00000143850 | PLEKHA6 | protein_coding | 0,664 | 1.301e-05 | 2.861e-03 | 782,9 | 496,71 | 821,847 | 768,176 | 758,668 | 490,707 | 551,618 | 447,803 |
| ENSG00000284341 | ENSG00000284341 | protein_coding | 0,664 | 1.436e-04 | 1.655e-02 | 8298,07 | 5239,97 | 8620,87 | 8595,537 | 7677,793 | 6251,403 | 5743,879 | 3724,622 |
| ENSG00000070495 | JMJD6 | protein_coding | 0,657 | 6.685e-08 | 3.326e-05 | 2438,57 | 1547,07 | 2436,189 | 2454,592 | 2424,935 | 1539,423 | 1545,812 | 1555,983 |
| ENSG00000177283 | FZD8 | protein_coding | 0,653 | 1.003e-04 | 1.317e-02 | 2097,97 | 1334,94 | 1801,814 | 2443,873 | 2048,229 | 1075,977 | 1423,943 | 1504,888 |

| | | | | | | | | | | | | | |
|---|---|---|---|---|---|---|---|---|---|---|---|---|---|
| ENSG00000270141 | TERC | lncRNA | 0,643 | 3.314e-04 | 3.042e-02 | 2052,58 | 1318,8 | 2021,478 | 2343,831 | 1792,419 | 1349,444 | 1611,557 | 995,386 |
| ENSG00000196872 | CRACDL | protein_coding | 0,64 | 9.464e-05 | 1.251e-02 | 1125,6 | 722,23 | 1004,585 | 1222,83 | 1149,391 | 587,826 | 772,906 | 805,949 |
| ENSG00000148297 | MED22 | protein_coding | 0,629 | 8.212e-05 | 1.126e-02 | 670,3 | 433,22 | 607,864 | 752,992 | 650,037 | 403,811 | 434,559 | 461,3 |
| ENSG00000145088 | EAF2 | protein_coding | 0,628 | 4.298e-04 | 3.645e-02 | 365,8 | 236,23 | 319,081 | 384,088 | 394,227 | 238,538 | 237,324 | 232,819 |
| ENSG00000051128 | HOMER3 | protein_coding | 0,613 | 2.109e-04 | 2.149e-02 | 5955,07 | 3897,02 | 5561,668 | 7574,577 | 4728,974 | 3802,127 | 4045,731 | 3843,2 |
| ENSG00000177469 | CAVIN1 | protein_coding | 0,61 | 8.856e-06 | 2.120e-03 | 4553,14 | 2982,63 | 4433,996 | 4948,486 | 4276,926 | 2632,439 | 3003,43 | 3312,007 |
| ENSG00000182253 | SYNM | protein_coding | 0,61 | 3.377e-05 | 5.883e-03 | 19376,99 | 12692,5 | 18744,356 | 20957,819 | 18428,805 | 11380,825 | 11399,564 | 15297,123 |
| ENSG00000184731 | FAM110C | protein_coding | 0,599 | 8.220e-05 | 1.126e-02 | 963,72 | 636,72 | 909,902 | 891,442 | 1089,819 | 639,793 | 684,711 | 585,663 |
| ENSG00000129474 | AJUBA | protein_coding | 0,594 | 5.042e-04 | 3.967e-02 | 1527,44 | 1014,74 | 1366,273 | 1915,975 | 1300,074 | 914,112 | 1109,65 | 1020,451 |
| ENSG00000197355 | UAP1L1 | protein_coding | 0,593 | 2.196e-04 | 2.209e-02 | 6041,04 | 4010,32 | 5847,61 | 7119,031 | 5156,491 | 3958,029 | 4667,904 | 3405,038 |
| ENSG00000130283 | GDF1 | protein_coding | 0,588 | 1.273e-04 | 1.523e-02 | 9265,89 | 6166,12 | 8204,266 | 11183,22 | 8410,179 | 5620,129 | 6930,499 | 5947,73 |
| ENSG00000148926 | ADM | protein_coding | 0,588 | 4.942e-04 | 3.943e-02 | 9478,33 | 6308,16 | 9171,924 | 9493,232 | 9769,825 | 6598,135 | 7706,612 | 4619,746 |
| ENSG00000248485 | PCP4L1 | protein_coding | 0,587 | 8.640e-05 | 1.167e-02 | 750,25 | 497,66 | 783,027 | 700,291 | 767,429 | 498,374 | 463,423 | 531,194 |
| ENSG00000184194 | GPR173 | protein_coding | 0,587 | 1.318e-04 | 1.567e-02 | 3174,05 | 2116,41 | 3226,79 | 3463,94 | 2831,427 | 2322,339 | 2297,872 | 1729,03 |
| ENSG00000065054 | NHERF2 | protein_coding | 0,585 | 3.869e-04 | 3.435e-02 | 751,74 | 504,71 | 763,143 | 828,023 | 664,054 | 478,78 | 583,688 | 451,659 |
| ENSG00000162729 | IGSF8 | protein_coding | 0,582 | 1.149e-04 | 1.421e-02 | 6384,14 | 4268,49 | 6280,311 | 7528,129 | 5343,968 | 4298,798 | 4637,437 | 3869,23 |
| ENSG00000223802 | CERS1 | protein_coding | 0,581 | 1.336e-04 | 1.579e-02 | 10558,57 | 7059,97 | 9566,751 | 12843,732 | 9265,214 | 6571,726 | 7749,908 | 6858,279 |
| ENSG00000113645 | WWC1 | protein_coding | 0,58 | 5.541e-06 | 1.397e-03 | 3633,78 | 2430,12 | 3662,331 | 3880,184 | 3358,815 | 2375,158 | 2331,547 | 2583,664 |
| ENSG00000185340 | GAS2L1 | protein_coding | 0,573 | 2.468e-04 | 2.431e-02 | 7081,06 | 4761,7 | 6339,961 | 8642,878 | 6260,327 | 4345,654 | 5341,391 | 4598,054 |
| ENSG00000174917 | MICOS13 | protein_coding | 0,573 | 5.979e-04 | 4.451e-02 | 1425,98 | 961,96 | 1286,739 | 1703,387 | 1287,809 | 845,958 | 1124,081 | 915,851 |
| ENSG00000178573 | MAF | protein_coding | 0,572 | 1.950e-06 | 6.047e-04 | 15001,47 | 10092,99 | 14274,381 | 15931,622 | 14798,411 | 10341,48 | 9497,766 | 10439,738 |
| ENSG00000139637 | MYG1 | protein_coding | 0,57 | 2.943e-04 | 2.755e-02 | 516,54 | 349,01 | 514,128 | 546,656 | 488,842 | 338,213 | 360,796 | 348,023 |
| ENSG00000142634 | EFHD2 | protein_coding | 0,559 | 6.397e-05 | 9.304e-03 | 6512,75 | 4422,86 | 6136,393 | 7560,285 | 5841,57 | 4588,451 | 4587,727 | 4092,408 |
| ENSG00000075043 | KCNQ2 | protein_coding | 0,55 | 1.459e-04 | 1.665e-02 | 6359,17 | 4344,96 | 5520,008 | 7134,216 | 6423,274 | 3963,993 | 4919,66 | 4151,216 |
| ENSG00000142627 | EPHA2 | protein_coding | 0,548 | 4.641e-05 | 7.324e-03 | 15317,61 | 10483,39 | 14410,724 | 16750,713 | 14791,402 | 9929,15 | 11869,401 | 9651,624 |
| ENSG00000139318 | DUSP6 | protein_coding | 0,538 | 4.412e-04 | 3.725e-02 | 22497,81 | 15498,92 | 22501,37 | 22664,778 | 22327,273 | 17688,455 | 16832,357 | 11975,958 |
| ENSG00000135404 | CD63 | protein_coding | 0,531 | 2.057e-04 | 2.137e-02 | 23357,92 | 16161,33 | 22505,157 | 27135,387 | 20433,231 | 15289,443 | 15249,663 | 17944,897 |
| ENSG00000127241 | MASP1 | protein_coding | 0,524 | 1.425e-04 | 1.653e-02 | 2037,77 | 1417,01 | 1979,818 | 1882,032 | 2251,475 | 1317,923 | 1531,38 | 1401,734 |
| ENSG00000113140 | SPARC | protein_coding | 0,514 | 1.888e-04 | 2.005e-02 | 60614,66 | 42437,9 | 59200,428 | 59409,517 | 63234,034 | 39192,667 | 38164,246 | 49956,784 |
| ENSG00000162706 | CADM3 | protein_coding | 0,513 | 8.862e-05 | 1.188e-02 | 1581,53 | 1109,11 | 1666,417 | 1534,566 | 1543,618 | 1156,91 | 1108,046 | 1062,387 |
| ENSG00000177875 | CCDC184 | protein_coding | 0,508 | 4.522e-05 | 7.246e-03 | 4726,87 | 3327,37 | 4620,521 | 5086,936 | 4473,164 | 3227,932 | 3510,148 | 3244,041 |
| ENSG00000070808 | CAMK2A | protein_coding | 0,508 | 2.706e-04 | 2.597e-02 | 4539,74 | 3191,84 | 4308,068 | 5079,79 | 4231,371 | 3015,804 | 2972,963 | 3586,762 |
| ENSG00000133985 | TTC9 | protein_coding | 0,505 | 2.738e-04 | 2.608e-02 | 3516,58 | 2474,18 | 3706,832 | 3419,278 | 3423,644 | 2499,539 | 2135,915 | 2787,079 |
| ENSG00000119771 | KLHL29 | protein_coding | 0,505 | 3.750e-04 | 3.361e-02 | 1394,62 | 983,53 | 1363,432 | 1569,402 | 1251,014 | 972,043 | 965,331 | 1013,221 |
| ENSG00000115457 | IGFBP2 | protein_coding | 0,487 | 6.238e-04 | 4.590e-02 | 64752,74 | 46216,5 | 59041,361 | 68073,832 | 67143,015 | 39670,596 | 53757,066 | 45221,834 |
| ENSG00000115310 | RTN4 | protein_coding | 0,479 | 1.561e-04 | 1.747e-02 | 31074,48 | 22294,63 | 29880,948 | 32170,515 | 31171,978 | 21286,121 | 20611,9 | 24985,864 |
| ENSG00000025039 | RRAGD | protein_coding | 0,468 | 2.007e-04 | 2.097e-02 | 9667,05 | 6985,02 | 9313,948 | 8908,167 | 10779,046 | 7083,731 | 6941,724 | 6929,619 |
| ENSG00000120049 | KCNIP2 | protein_coding | 0,464 | 3.702e-04 | 3.349e-02 | 4811,39 | 3492,23 | 4825,036 | 4994,04 | 4615,086 | 3363,388 | 3907,826 | 3205,479 |
| ENSG00000130821 | SLC6A8 | protein_coding | 0,436 | 6.681e-04 | 4.822e-02 | 10120,77 | 7482,17 | 9505,207 | 10293,565 | 10563,535 | 6736,147 | 8174,846 | 7535,526 |
| ENSG00000130558 | OLFM1 | protein_coding | 0,424 | 4.276e-04 | 3.645e-02 | 5448,55 | 4062,45 | 5668,66 | 5446,907 | 5230,08 | 4156,527 | 4194,86 | 3835,97 |
| ENSG00000133056 | PIK3C2B | protein_coding | 0,402 | 5.099e-04 | 3.984e-02 | 13632,34 | 10314,19 | 13568,047 | 14280,936 | 13048,042 | 10273,326 | 10121,543 | 10547,712 |
| ENSG00000186638 | KIF24 | protein_coding | -0,442 | 2.690e-04 | 2.595e-02 | 1886,42 | 2562,99 | 1924,902 | 1854,342 | 1880,025 | 2618,808 | 2540,007 | 2530,159 |
| ENSG00000163808 | KIF15 | protein_coding | -0,443 | 2.863e-04 | 2.693e-02 | 2393,47 | 3254,61 | 2339,612 | 2491,214 | 2349,594 | 3145,296 | 3208,683 | 3409,858 |
| ENSG00000138434 | ITPRID2 | protein_coding | -0,447 | 6.886e-04 | 4.932e-02 | 23952,05 | 32656,22 | 23712,363 | 22236,922 | 25906,856 | 30460,468 | 30757,496 | 36750,694 |
| ENSG00000130066 | SAT1 | protein_coding | -0,467 | 4.286e-04 | 3.645e-02 | 1903,85 | 2633,5 | 2117,108 | 1730,183 | 1864,256 | 2642,662 | 2693,947 | 2563,901 |
| ENSG00000038427 | VCAN | protein_coding | -0,471 | 2.628e-04 | 2.548e-02 | 27933,19 | 38719,59 | 29289,181 | 24529,839 | 29980,536 | 38729,222 | 36631,262 | 40798,274 |
| ENSG00000136492 | BRIP1 | protein_coding | -0,489 | 5.167e-04 | 3.990e-02 | 2466,32 | 3458,27 | 2573,479 | 2104,446 | 2721,043 | 3716,935 | 3304,895 | 3352,979 |
| ENSG00000117385 | P3H1 | protein_coding | -0,502 | 3.520e-04 | 3.216e-02 | 1251,22 | 1776,12 | 1221,408 | 1389,863 | 1142,383 | 1673,175 | 1929,058 | 1726,138 |
| ENSG00000048052 | HDAC9 | protein_coding | -0,514 | 1.993e-04 | 2.094e-02 | 4289,32 | 6119,55 | 4344,994 | 3725,656 | 4797,306 | 6027,348 | 5719,826 | 6611,481 |
| ENSG00000213988 | ZNF90 | protein_coding | -0,514 | 4.729e-04 | 3.862e-02 | 854,57 | 1217,72 | 927,892 | 740,486 | 895,334 | 1276,179 | 1144,927 | 1232,061 |
| ENSG00000103888 | CEMIP | protein_coding | -0,517 | 5.751e-04 | 4.333e-02 | 695,74 | 995,76 | 726,217 | 618,114 | 742,899 | 918,372 | 1090,407 | 978,515 |
| ENSG00000158402 | CDC25C | protein_coding | -0,523 | 3.947e-04 | 3.488e-02 | 510,49 | 732,41 | 525,49 | 469,838 | 536,149 | 713,911 | 747,25 | 736,055 |
| ENSG00000111331 | OAS3 | protein_coding | -0,523 | 4.286e-04 | 3.645e-02 | 594,14 | 856,28 | 632,481 | 575,239 | 574,696 | 888,555 | 907,603 | 772,689 |
| ENSG00000214548 | MEG3 | lncRNA | -0,527 | 5.167e-04 | 3.990e-02 | 747,47 | 1081,36 | 847,411 | 744,953 | 650,037 | 1051,272 | 1164,17 | 1028,645 |
| ENSG00000258830 | ENSG00000258830 | protein_coding | -0,545 | 6.207e-04 | 4.585e-02 | 1611,94 | 2346,86 | 1676,833 | 1317,512 | 1841,479 | 2255,889 | 2116,672 | 2668,019 |
| ENSG00000106804 | C5 | protein_coding | -0,547 | 3.008e-04 | 2.802e-02 | 1137,83 | 1658,38 | 1077,49 | 979,872 | 1356,141 | 1677,434 | 1685,32 | 1612,38 |
| ENSG00000283201 | ENSG00000283201 | protein_coding | -0,554 | 1.213e-04 | 1.470e-02 | 967,52 | 1415,86 | 994,17 | 883,403 | 1024,991 | 1478,936 | 1255,572 | 1513,082 |
| ENSG00000008311 | AASS | protein_coding | -0,558 | 2.374e-05 | 4.581e-03 | 3289,63 | 4843,26 | 3447,401 | 3371,044 | 3050,442 | 5282,768 | 4342,386 | 4904,624 |
| ENSG00000000460 | FIRRM | protein_coding | -0,558 | 1.483e-04 | 1.679e-02 | 915,2 | 1342,4 | 865,401 | 828,916 | 1051,272 | 1289,81 | 1305,281 | 1432,102 |
| ENSG00000123572 | NRK | protein_coding | -0,558 | 1.663e-04 | 1.817e-02 | 598,08 | 877,39 | 604,076 | 571,666 | 618,499 | 868,108 | 798,563 | 965,5 |
| ENSG00000158258 | CLSTN2 | protein_coding | -0,562 | 1.223e-04 | 1.473e-02 | 6910,73 | 10197,59 | 6549,21 | 5856,899 | 8326,077 | 10086,755 | 10196,909 | 10309,109 |
| ENSG00000125354 | SEPTIN6 | protein_coding | -0,565 | 2.346e-04 | 2.335e-02 | 477,72 | 704,56 | 485,723 | 432,323 | 515,123 | 659,388 | 716,782 | 737,501 |
| ENSG00000270647 | TAF15 | protein_coding | -0,565 | 3.237e-04 | 2.986e-02 | 2372,61 | 3507,31 | 2530,872 | 2274,16 | 2312,799 | 3515,03 | 2809,402 | 4197,49 |
| ENSG00000091513 | TF | protein_coding | -0,575 | 5.170e-04 | 3.990e-02 | 538,18 | 803,85 | 601,236 | 485,916 | 527,388 | 700,28 | 928,45 | 782,812 |
| ENSG00000151725 | CENPU | protein_coding | -0,576 | 2.100e-04 | 2.149e-02 | 1278,77 | 1901,01 | 1243,185 | 1184,421 | 1408,705 | 1678,286 | 1765,497 | 2259,26 |
| ENSG00000073910 | FRY | protein_coding | -0,58 | 2.161e-04 | 2.185e-02 | 466,41 | 693,62 | 465,839 | 430,536 | 502,859 | 675,574 | 639,812 | 765,459 |
| ENSG00000137033 | IL33 | protein_coding | -0,58 | 5.029e-04 | 3.967e-02 | 671,15 | 999,18 | 625,853 | 588,638 | 798,967 | 1140,723 | 938,071 | 918,743 |
| ENSG00000168743 | NPNT | protein_coding | -0,583 | 4.537e-04 | 3.779e-02 | 8058,36 | 12063,13 | 7477,102 | 6896,617 | 9801,363 | 10709,51 | 11005,093 | 14474,785 |
| ENSG00000164684 | ZNF704 | protein_coding | -0,584 | 4.870e-06 | 1.297e-03 | 7885,62 | 11812,13 | 7823,641 | 7119,924 | 8713,296 | 11920,943 | 11248,831 | 12266,62 |
| ENSG00000155849 | ELMO1 | protein_coding | -0,584 | 4.452e-04 | 3.742e-02 | 414,35 | 619,04 | 458,265 | 355,505 | 429,27 | 668,759 | 566,05 | 622,297 |
| ENSG00000144339 | TMEFF2 | protein_coding | -0,589 | 1.507e-04 | 1.696e-02 | 525,64 | 791,03 | 467,733 | 597,57 | 511,619 | 767,582 | 779,32 | 826,194 |
| ENSG00000137145 | DENND4C | protein_coding | -0,6 | 1.686e-05 | 3.543e-03 | 3743,99 | 5670,15 | 3684,108 | 3404,094 | 4143,765 | 5485,525 | 5136,138 | 6388,784 |
| ENSG00000117602 | RCAN3 | protein_coding | -0,603 | 3.724e-04 | 3.353e-02 | 862,24 | 1303,52 | 764,09 | 843,208 | 979,435 | 1218,248 | 1111,253 | 1581,048 |
| ENSG00000162631 | NTNG1 | protein_coding | -0,605 | 5.492e-04 | 4.169e-02 | 242,43 | 366,82 | 248,069 | 228,666 | 250,553 | 375,698 | 336,743 | 388,032 |
| ENSG00000156103 | MMP16 | protein_coding | -0,607 | 4.580e-04 | 3.799e-02 | 19637,87 | 29901,76 | 19260,377 | 16460,414 | 23192,821 | 26323,535 | 25693,517 | 37688,236 |
| ENSG00000138658 | ZGRF1 | protein_coding | -0,61 | 1.335e-05 | 2.902e-03 | 1634,38 | 2493,05 | 1695,769 | 1418,447 | 1788,915 | 2583,879 | 2580,095 | 2315,175 |
| ENSG00000183850 | ZNF730 | protein_coding | -0,611 | 2.981e-05 | 5.420e-03 | 1021,4 | 1554,54 | 943,041 | 968,26 | 1152,895 | 1662,1 | 1393,476 | 1608,042 |
| ENSG00000079689 | SCGN | protein_coding | -0,611 | 4.855e-04 | 3.907e-02 | 688,97 | 1049,97 | 623,013 | 783,361 | 660,549 | 989,933 | 873,929 | 1286,048 |
| ENSG00000243244 | STON1 | protein_coding | -0,612 | 6.058e-04 | 4.492e-02 | 349,12 | 529,99 | 341,805 | 335,854 | 369,697 | 480,484 | 469,837 | 639,65 |

| | | | | | | | | | | | | | |
|---|---|---|---|---|---|---|---|---|---|---|---|---|---|
| ENSG00000163541 | SUCLG1 | protein_coding | -0,614 | 5.473e-04 | 4.169e-02 | 2268,94 | 3466,79 | 2199,482 | 1830,225 | 2777,111 | 3020,915 | 3147,749 | 4231,714 |
| ENSG00000134121 | CHL1 | protein_coding | -0,618 | 1.124e-04 | 1.402e-02 | 5387,74 | 8264,19 | 5196,193 | 4447,384 | 6519,641 | 7926,281 | 7448,442 | 9417,841 |
| ENSG00000114861 | FOXP1 | protein_coding | -0,618 | 4.110e-04 | 3.594e-02 | 533,88 | 814,94 | 582,299 | 435,895 | 583,456 | 791,435 | 711,972 | 941,399 |
| ENSG00000152527 | PLEKHH2 | protein_coding | -0,623 | 1.522e-05 | 3.270e-03 | 2091,33 | 3221,68 | 2341,506 | 1907,043 | 2025,451 | 3653,893 | 3061,157 | 2950,005 |
| ENSG00000260804 | LINC01963 | lncRNA | -0,624 | 2.084e-04 | 2.149e-02 | 2838,87 | 4368,56 | 2928,539 | 2626,985 | 2961,084 | 4214,458 | 3399,504 | 5491,732 |
| ENSG00000128833 | MYO5C | protein_coding | -0,626 | 1.062e-05 | 2.449e-03 | 6566,81 | 10126,91 | 6802,013 | 5844,394 | 7054,038 | 9625,866 | 9088,863 | 11666,015 |
| ENSG00000206190 | ATP10A | protein_coding | -0,627 | 1.473e-06 | 4.801e-04 | 1268,88 | 1959,66 | 1203,419 | 1345,202 | 1258,023 | 1868,265 | 1961,129 | 2049,578 |
| ENSG00000150471 | ADGRL3 | protein_coding | -0,63 | 3.139e-06 | 8.602e-04 | 1428,84 | 2210,72 | 1491,254 | 1446,137 | 1349,133 | 2143,436 | 2042,91 | 2445,804 |
| ENSG00000013619 | MAMLD1 | protein_coding | -0,639 | 4.428e-06 | 1.196e-03 | 747,33 | 1163,62 | 799,123 | 729,768 | 713,113 | 1131,352 | 1124,081 | 1235,435 |
| ENSG00000021826 | CPS1 | protein_coding | -0,64 | 1.033e-04 | 1.347e-02 | 505,34 | 785,42 | 492,351 | 433,216 | 590,465 | 804,214 | 838,651 | 713,4 |
| ENSG00000140092 | FBLN5 | protein_coding | -0,642 | 1.631e-04 | 1.793e-02 | 327,51 | 509,81 | 345,592 | 276,008 | 360,937 | 508,597 | 529,168 | 491,667 |
| ENSG00000121966 | CXCR4 | protein_coding | -0,645 | 1.390e-04 | 1.632e-02 | 1350,97 | 2106,21 | 1200,578 | 1179,062 | 1673,275 | 1863,153 | 2028,478 | 2427,005 |
| ENSG00000157510 | AFAP1L1 | protein_coding | -0,645 | 2.114e-04 | 2.149e-02 | 1086,09 | 1697,29 | 1120,098 | 850,353 | 1287,809 | 1668,915 | 1954,715 | 1468,254 |
| ENSG00000101938 | CHRDL1 | protein_coding | -0,655 | 1.172e-05 | 2.638e-03 | 1445,21 | 2278,22 | 1617,182 | 1369,319 | 1349,133 | 2576,212 | 2285,044 | 1973,418 |
| ENSG00000305512 | ENSG00000305512 | lncRNA | -0,659 | 1.264e-05 | 2.812e-03 | 396,84 | 624,88 | 392,934 | 384,088 | 413,5 | 620,199 | 610,949 | 643,506 |
| ENSG00000290217 | ENSG00000290217 | protein_coding | -0,667 | 2.745e-04 | 2.608e-02 | 431,69 | 683,72 | 433,647 | 468,945 | 392,475 | 653,424 | 553,221 | 844,511 |
| ENSG00000140750 | ARHGAP17 | protein_coding | -0,669 | 2.012e-06 | 6.047e-04 | 1645,97 | 2614,82 | 1514,925 | 1697,134 | 1725,839 | 2274,631 | 2724,414 | 2845,405 |
| ENSG00000281593 | ENSG00000281593 | protein_coding | -0,674 | 8.351e-05 | 1.136e-02 | 350,93 | 557,01 | 375,891 | 303,698 | 373,202 | 531,599 | 514,736 | 624,707 |
| ENSG00000205413 | SAMD9 | protein_coding | -0,681 | 6.654e-05 | 9.539e-03 | 446,61 | 714,1 | 444,062 | 478,77 | 417,005 | 723,282 | 588,499 | 830,532 |
| ENSG00000174640 | SLCO2A1 | protein_coding | -0,691 | 4.739e-04 | 3.862e-02 | 312,36 | 500,44 | 283,102 | 250,997 | 402,988 | 441,296 | 540,393 | 519,625 |
| ENSG00000131373 | HACL1 | protein_coding | -0,694 | 1.046e-04 | 1.354e-02 | 318,91 | 512,08 | 320,028 | 263,502 | 373,202 | 475,372 | 509,926 | 550,957 |
| ENSG00000120071 | KANSL1 | protein_coding | -0,703 | 4.799e-04 | 3.878e-02 | 1005,96 | 1632,48 | 1139,034 | 713,69 | 1165,16 | 1415,894 | 1427,15 | 2054,399 |
| ENSG00000079102 | RUNX1T1 | protein_coding | -0,705 | 4.919e-04 | 3.941e-02 | 181,61 | 295,96 | 212,089 | 173,286 | 159,443 | 299,025 | 258,17 | 330,67 |
| ENSG00000290683 | ENSG00000290683 | lncRNA | -0,713 | 9.569e-08 | 4.523e-05 | 1003,48 | 1645,05 | 1028,255 | 1030,786 | 951,402 | 1757,515 | 1547,416 | 1630,215 |
| ENSG00000105339 | DENND3 | protein_coding | -0,721 | 1.068e-04 | 1.355e-02 | 186,58 | 306,48 | 190,312 | 167,927 | 201,494 | 302,432 | 314,294 | 302,713 |
| ENSG00000151388 | ADAMTS12 | protein_coding | -0,722 | 5.422e-06 | 1.385e-03 | 348,03 | 573,14 | 337,071 | 367,117 | 339,911 | 597,197 | 525,961 | 596,267 |
| ENSG00000106144 | CASP2 | protein_coding | -0,73 | 2.933e-07 | 1.206e-04 | 2579,47 | 4278,66 | 2797,877 | 2575,178 | 2365,363 | 4274,944 | 3718,609 | 4842,442 |
| ENSG00000203497 | PDCD4-AS1 | lncRNA | -0,737 | 1.616e-04 | 1.787e-02 | 160,37 | 266,18 | 160,014 | 159,888 | 161,195 | 253,873 | 250,152 | 294,518 |
| ENSG00000134874 | DZIP1 | protein_coding | -0,741 | 1.569e-10 | 1.484e-07 | 3608,66 | 6029,79 | 3621,618 | 3582,739 | 3621,633 | 5976,232 | 5952,34 | 6160,786 |
| ENSG00000127152 | BCL11B | protein_coding | -0,745 | 6.036e-06 | 1.502e-03 | 385,32 | 645,68 | 382,519 | 403,739 | 369,697 | 565,676 | 638,209 | 733,163 |
| ENSG00000185774 | KCNIP4 | protein_coding | -0,752 | 6.608e-04 | 4.787e-02 | 150,74 | 250,21 | 145,812 | 125,945 | 180,468 | 221,5 | 234,117 | 295 |
| ENSG00000221963 | APOL6 | protein_coding | -0,756 | 4.996e-04 | 3.967e-02 | 159,48 | 265,4 | 124,981 | 157,208 | 196,238 | 263,244 | 238,927 | 294,036 |
| ENSG00000303606 | ENSG00000303606 | lncRNA | -0,756 | 5.918e-04 | 4.441e-02 | 130,95 | 220,21 | 135,396 | 131,305 | 126,153 | 258,984 | 177,993 | 223,66 |
| ENSG00000157191 | NECAP2 | protein_coding | -0,758 | 2.157e-07 | 9.270e-05 | 687,4 | 1161,84 | 630,588 | 736,02 | 695,592 | 1064,05 | 1160,963 | 1260,5 |
| ENSG00000173406 | DAB1 | protein_coding | -0,758 | 9.033e-06 | 2.135e-03 | 551,84 | 932,09 | 607,864 | 450,187 | 597,473 | 1035,085 | 950,899 | 810,287 |
| ENSG00000107438 | PDLIM1 | protein_coding | -0,761 | 4.128e-07 | 1.626e-04 | 888,6 | 1504,04 | 953,456 | 755,671 | 956,658 | 1616,096 | 1512,138 | 1383,899 |
| ENSG00000154856 | APCDD1 | protein_coding | -0,764 | 6.786e-07 | 2.516e-04 | 697,46 | 1178,48 | 672,248 | 640,445 | 779,694 | 1155,206 | 1055,129 | 1325,092 |
| ENSG00000272398 | CD24 | protein_coding | -0,765 | 3.207e-05 | 5.775e-03 | 247,87 | 419,25 | 249,963 | 250,104 | 243,545 | 396,144 | 365,607 | 496,006 |
| ENSG00000149548 | CCDC15 | protein_coding | -0,77 | 2.047e-06 | 6.047e-04 | 687,16 | 1171,52 | 737,579 | 602,036 | 721,874 | 1351,148 | 1159,359 | 1004,062 |
| ENSG00000140553 | UNC45A | protein_coding | -0,773 | 9.517e-07 | 3.381e-04 | 1700,16 | 2909,59 | 1549,011 | 1955,277 | 1596,182 | 2621,364 | 3462,042 | 2645,364 |
| ENSG00000110811 | P3H3 | protein_coding | -0,787 | 2.456e-06 | 6.908e-04 | 815,53 | 1413,68 | 799,123 | 936,103 | 711,361 | 1347,74 | 1680,51 | 1212,78 |
| ENSG00000152954 | NRSN1 | protein_coding | -0,794 | 4.264e-04 | 3.645e-02 | 124,98 | 217,47 | 136,343 | 107,187 | 131,409 | 232,575 | 240,531 | 179,314 |
| ENSG00000128602 | SMO | protein_coding | -0,799 | 2.625e-09 | 2.068e-06 | 2472,59 | 4304,26 | 2330,144 | 2660,928 | 2426,687 | 3855,799 | 4724,028 | 4332,94 |
| ENSG00000164946 | FREM1 | protein_coding | -0,803 | 4.388e-08 | 2.242e-05 | 514,08 | 895,44 | 518,862 | 508,247 | 515,123 | 970,339 | 819,409 | 896,57 |
| ENSG00000144290 | SLC4A10 | protein_coding | -0,808 | 9.833e-07 | 3.381e-04 | 254,06 | 445,2 | 249,016 | 262,609 | 250,553 | 442,999 | 450,595 | 442,019 |
| ENSG00000283982 | ENSG00000283982 | lncRNA | -0,811 | 4.221e-04 | 3.645e-02 | 102,02 | 180,35 | 119,3 | 99,148 | 87,606 | 193,386 | 173,182 | 174,494 |
| ENSG00000137275 | RIPK1 | protein_coding | -0,812 | 4.670e-09 | 3.271e-06 | 1415,05 | 2484,16 | 1561,32 | 1390,757 | 1293,065 | 2530,208 | 2260,991 | 2661,27 |
| ENSG00000143494 | VASH2 | protein_coding | -0,817 | 8.688e-08 | 4.212e-05 | 454,84 | 797,08 | 422,285 | 434,109 | 508,115 | 792,287 | 761,681 | 837,281 |
| ENSG00000146833 | TRIM4 | protein_coding | -0,82 | 2.919e-09 | 2.208e-06 | 876,82 | 1552,29 | 873,922 | 959,327 | 797,215 | 1501,086 | 1619,575 | 1536,22 |
| ENSG00000151572 | ANO4 | protein_coding | -0,832 | 2.484e-06 | 6.908e-04 | 1144,32 | 2030,34 | 1057,607 | 975,405 | 1399,944 | 1783,924 | 1837,657 | 2469,424 |
| ENSG00000137868 | STRA6 | protein_coding | -0,853 | 6.070e-07 | 2.342e-04 | 529,7 | 954,71 | 463,946 | 517,179 | 607,986 | 789,732 | 1077,579 | 996,832 |
| ENSG00000141622 | ARK2C | protein_coding | -0,855 | 1.184e-07 | 5.459e-05 | 1877,34 | 3394,47 | 1902,178 | 1572,975 | 2156,861 | 3416,207 | 3907,826 | 2859,383 |
| ENSG00000111058 | ACSS3 | protein_coding | -0,857 | 2.851e-08 | 1.685e-05 | 2464,33 | 4456,04 | 2395,475 | 2211,634 | 2785,872 | 4069,631 | 3988,003 | 5310,49 |
| ENSG00000196569 | LAMA2 | protein_coding | -0,862 | 1.165e-04 | 1.431e-02 | 108,8 | 198,64 | 107,938 | 108,081 | 110,384 | 215,536 | 211,667 | 168,709 |
| ENSG00000272970 | ENSG00000272970 | lncRNA | -0,874 | 2.077e-05 | 4.160e-03 | 271,1 | 493,35 | 269,846 | 198,297 | 345,168 | 532,451 | 509,926 | 437,68 |
| ENSG00000230797 | YY2 | protein_coding | -0,893 | 9.351e-05 | 1.245e-02 | 136,86 | 253,18 | 147,705 | 140,237 | 122,648 | 288,802 | 184,407 | 286,324 |
| ENSG00000275713 | H2BC9 | protein_coding | -0,9 | 4.687e-05 | 7.324e-03 | 1259,9 | 2345,17 | 1212,887 | 909,307 | 1657,506 | 1649,321 | 2390,878 | 2995,315 |
| ENSG00000197635 | DPP4 | protein_coding | -0,905 | 4.964e-06 | 1.304e-03 | 405,65 | 755,97 | 415,658 | 394,807 | 406,492 | 663,647 | 594,913 | 1009,364 |
| ENSG00000065361 | ERBB3 | protein_coding | -0,92 | 3.589e-08 | 1.996e-05 | 556,19 | 1053,81 | 563,363 | 527,005 | 578,2 | 1165,429 | 1157,756 | 838,245 |
| ENSG00000116106 | EPHA4 | protein_coding | -0,926 | 7.217e-11 | 8.027e-08 | 730,49 | 1385,63 | 778,293 | 642,231 | 770,933 | 1398,004 | 1396,683 | 1362,208 |
| ENSG00000106785 | TRIM14 | protein_coding | -0,958 | 6.691e-05 | 9.539e-03 | 214,89 | 411,77 | 160,014 | 192,044 | 292,604 | 327,138 | 383,246 | 524,927 |
| ENSG00000143507 | DUSP10 | protein_coding | -0,969 | 3.106e-07 | 1.249e-04 | 177,67 | 345,67 | 183,685 | 170,607 | 178,716 | 351,844 | 301,465 | 383,693 |
| ENSG00000305782 | ENSG00000305782 | lncRNA | -0,969 | 2.103e-05 | 4.160e-03 | 182,82 | 353,15 | 179,897 | 123,266 | 245,297 | 336,509 | 362,4 | 360,556 |
| ENSG00000107518 | ATRNL1 | protein_coding | -0,985 | 3.838e-05 | 6.537e-03 | 93,6 | 183,15 | 74,799 | 97,362 | 108,631 | 166,125 | 182,804 | 200,523 |
| ENSG00000005471 | ABCB4 | protein_coding | -1,008 | 1.633e-06 | 5.235e-04 | 138,16 | 278,06 | 158,12 | 119,693 | 136,665 | 313,507 | 270,998 | 249,69 |
| ENSG00000156113 | KCNMA1 | protein_coding | -1,017 | 5.511e-04 | 4.169e-02 | 59,75 | 120,55 | 78,587 | 41,089 | 59,572 | 127,788 | 113,851 | 120,025 |
| ENSG00000188641 | DPYD | protein_coding | -1,026 | 4.017e-04 | 3.533e-02 | 53,06 | 108,31 | 54,916 | 46,448 | 57,82 | 92,859 | 129,887 | 102,19 |
| ENSG00000105357 | MYH14 | protein_coding | -1,038 | 2.230e-07 | 9.371e-05 | 1119,29 | 2304,82 | 1055,713 | 1543,499 | 758,668 | 2284,003 | 2273,819 | 2356,629 |
| ENSG00000114779 | ABHD14B | protein_coding | -1,062 | 4.442e-09 | 3.231e-06 | 209,28 | 439,71 | 191,259 | 243,851 | 192,733 | 436,184 | 460,216 | 422,738 |
| ENSG00000129219 | PLD2 | protein_coding | -1,074 | 2.770e-11 | 3.274e-08 | 1051,86 | 2216,32 | 1255,494 | 906,627 | 993,452 | 1911,713 | 2374,842 | 2362,414 |
| ENSG00000113805 | CNTN3 | protein_coding | -1,089 | 4.292e-05 | 7.057e-03 | 69,39 | 146,89 | 61,544 | 62,526 | 84,102 | 140,567 | 174,786 | 125,327 |
| ENSG00000074621 | SLC24A1 | protein_coding | -1,092 | 1.647e-14 | 2.396e-11 | 567,29 | 1205,9 | 534,958 | 546,656 | 620,251 | 1264,252 | 1185,016 | 1168,433 |
| ENSG00000185070 | FLRT2 | protein_coding | -1,151 | 1.462e-04 | 1.665e-02 | 43,36 | 94,69 | 42,607 | 38,409 | 49,059 | 101,379 | 83,384 | 99,298 |
| ENSG00000183780 | SLC35F3 | protein_coding | -1,169 | 3.892e-05 | 6.570e-03 | 53,73 | 121,12 | 57,757 | 49,128 | 54,316 | 96,267 | 137,904 | 129,183 |
| ENSG00000136068 | FLNB | protein_coding | -1,186 | 7.139e-19 | 1.350e-15 | 1308,65 | 2975,87 | 1368,167 | 1259,452 | 1298,321 | 2767,894 | 2860,715 | 3298,992 |
| ENSG00000172137 | CALB2 | protein_coding | -1,242 | 1.572e-04 | 1.748e-02 | 70,66 | 165,95 | 58,703 | 90,216 | 63,076 | 129,492 | 109,041 | 259,33 |
| ENSG00000238083 | LRRC37A2 | protein_coding | -1,256 | 5.087e-06 | 1.318e-03 | 50,96 | 121,09 | 53,022 | 45,555 | 54,316 | 127,788 | 120,265 | 115,204 |
| ENSG00000100889 | PCK2 | protein_coding | -1,356 | 1.405e-04 | 1.640e-02 | 35,54 | 91,23 | 35,979 | 28,583 | 42,051 | 64,746 | 129,887 | 79,052 |
| ENSG00000166147 | FBN1 | protein_coding | -1,37 | 7.096e-24 | 1.917e-20 | 1055,83 | 2729,82 | 1152,29 | 1016,494 | 998,709 | 2727,854 | 2520,764 | 2940,846 |

| | | | | | | | | | | | | | |
|---|---|---|---|---|---|---|---|---|---|---|---|---|---|
| ENSG00000186854 | TRABD2A | protein_coding | -1,399 | 2.330e-04 | 2.331e-02 | 24,7 | 64,57 | 25,564 | 18,758 | 29,786 | 66,45 | 73,763 | 53,505 |
| ENSG00000168528 | SERINC2 | protein_coding | -1,458 | 6.430e-09 | 4.342e-06 | 91,37 | 251,68 | 109,832 | 69,672 | 94,615 | 283,69 | 279,016 | 192,329 |
| ENSG00000161791 | FMNL3 | protein_coding | -1,469 | 5.074e-25 | 1.599e-21 | 1569,21 | 4345,3 | 1645,587 | 1651,579 | 1410,457 | 4188,9 | 3906,223 | 4940,776 |
| ENSG00000123427 | EEF1AKMT3 | protein_coding | -1,562 | 6.776e-15 | 1.068e-11 | 132,76 | 388,96 | 130,662 | 113,44 | 154,187 | 343,325 | 429,749 | 393,816 |
| ENSG00000166535 | A2ML1 | protein_coding | -1,604 | 1.318e-07 | 5.934e-05 | 40,29 | 123,16 | 49,235 | 34,836 | 36,795 | 149,086 | 113,851 | 106,528 |
| ENSG00000140575 | IQGAP1 | protein_coding | -1,633 | 2.084e-23 | 4.378e-20 | 4301,51 | 13331,07 | 4515,423 | 3553,263 | 4835,853 | 12000,172 | 11784,414 | 16208,636 |
| ENSG00000011304 | PTBP1 | protein_coding | -1,731 | 1.594e-34 | 6.029e-31 | 4962,33 | 16468,45 | 4754,024 | 5174,472 | 4958,501 | 14050,748 | 16168,491 | 19186,116 |
| ENSG00000081923 | ATP8B1 | protein_coding | -1,791 | 6.116e-05 | 9.035e-03 | 14,51 | 49,3 | 17,043 | 10,719 | 15,769 | 53,671 | 41,692 | 52,541 |
| ENSG00000185559 | DLK1 | protein_coding | -1,832 | 9.676e-11 | 1.016e-07 | 50,1 | 180,82 | 48,288 | 65,206 | 36,795 | 161,013 | 158,75 | 222,696 |
| ENSG00000122786 | CALD1 | protein_coding | -2,226 | 6.196e-65 | 3.905e-61 | 3475,31 | 16252,59 | 3493,796 | 3366,578 | 3565,566 | 15084,129 | 15195,143 | 18478,501 |
| ENSG00000258744 | ENSG00000258744 | lncRNA | -2,288 | 1.522e-23 | 3.598e-20 | 73,13 | 356 | 66,278 | 81,284 | 71,837 | 415,738 | 280,619 | 371,643 |
| ENSG00000196924 | FLNA | protein_coding | -2,376 | 1.720e-82 | 3.253e-78 | 25494,52 | 132375,3 | 25835,152 | 25563,305 | 25085,111 | 118770,693 | 144052,389 | 134302,813 |
